# Targeting peptide–MHC complexes with designed T cell receptors and antibodies

**DOI:** 10.1101/2025.11.19.689381

**Authors:** Amir Motmaen, Kevin M. Jude, Nan Wang, Anastasia Minervina, David Feldman, Mauriz A. Lichtenstein, Abishai Ebenezer, Colin Correnti, Paul G. Thomas, K. Christopher Garcia, David Baker, Philip Bradley

**Author notes:** Corresponding author. (K.C.G), (D.B.), (P.B.). These authors contributed equally to this work (Kevin M. Jude, Nan Wang).

## Abstract

Class I major histocompatibility complexes (MHCs), expressed on the surface of all nucleated cells, present peptides derived from intracellular proteins for surveillance by T cells. The precise recognition of foreign or mutated peptide–MHC (pMHC) complexes by T cell receptors (TCRs) is central to immune defense against pathogens and tumors. Although patient-derived TCRs specific for cancer-associated antigens have been used to engineer tumor-targeting therapies, their reactivity toward self- or near-self antigens may be constrained by negative selection in the thymus. Here, we introduce a structure-based deep learning framework, ADAPT (Antigen-receptor Design Against Peptide-MHC Targets), for the design of TCRs and antibodies that bind to pMHC targets of interest. We evaluate the ADAPT pipeline by designing and characterizing TCRs and antibodies against a diverse panel of pMHCs. Cryogenic electron microscopy structures of two designed antibodies bound to their respective pMHC targets demonstrate atomic-level accuracy at the recognition interface, supporting the robustness of our structure-based approach. Computationally designed TCRs and antibodies targeting pMHC complexes could enable a broad range of therapeutic applications, from cancer immunotherapy to autoimmune disease treatment, and insights gained from TCR–pMHC design should advance predictive understanding of TCR specificity with implications for basic immunology and clinical diagnostics.

## Introduction

The peptide–MHC complexes (pMHCs) on the surface of a cell provide a window onto its internally expressed proteome, enabling the detection of viral infection or oncogenic transformation from external cues. Thus disease-associated pMHCs are an attractive recognition target for delivery of cytotoxic therapies, and this is the basis of the cellular arm of the adaptive immune system, in which T cell receptors (TCRs) mediate recognition (*1, 2*). Highly variable TCRs are produced by a genome rearrangement process—V(D)J recombination—that generates sequence diversity by random joining of germline-encoded gene segments together with nucleotide insertion and deletion at the segment junctions (*3*). TCR sequences are filtered during T cell development in the thymus by positive and negative selection processes that guarantee a baseline but not excessively strong level of binding to self-pMHCs (*4*). Native TCRs derived from patients are being explored as therapeutics, but their reactivity toward self or near-self targets may be constrained by thymic selection. Computational protein design represents an alternative path to therapeutic TCRs, but de novo generation of the loop-mediated binding modes employed by TCRs remains challenging, and past work on TCR design has been limited to the reengineering of existing binders (*5–9*). The composite nature of the pMHC interface—consisting of a few variable peptide residues surrounded by many conserved MHC positions—represents an additional barrier to peptide-specific recognition.

Deep learning protein design methods based on generative diffusion models (*10*) or gradient backpropagation through structure prediction networks (*11*) have been widely used to generate binders against many targets. Though successful, these methods can be compute- and memory-intensive for large systems like TCR:pMHC complexes, and network backpropagation may lead to inflated model confidence and exploration of adversarial sequences and structures in some cases. We reasoned that the highly conserved binding modes of TCRs for their pMHC targets would be well-learned by structure prediction networks trained on the PDB and fine-tuned on TCRs, enabling the use of simpler inference-based methods directly for conformational sampling. We set out to explore the possibility of combining such an approach for conformational sampling with ProteinMPNN-based sequence design (*12*) to generate pMHC-specific TCRs and ‘TCR-mimic’ antibodies.

## Results

We developed a structure-based deep learning strategy, ADAPT (Antigen-receptor Design Against Peptide-MHC Targets), for the design of pMHC-targeting TCRs and antibodies. The ADAPT design pipeline (**Fig. 1**) combines fine-tuned Alphafold2 (*13*) (AF2) conformational sampling with ProteinMPNN (*12*) fixed-backbone sequence design. Diversity is introduced into the pipeline through the use of multiple TCR/antibody template structures (the choice of template determines the sequence and structure of the receptor outside the Complementarity-Determining Region (CDR) loops) and by initializing each design run with random CDR3 sequences taken from a large library of paired human TCRs. A single design run consists of four steps (**Fig. 1**, top): initial docking, sequence design of the 6 CDR loops, redocking, and design model evaluation. TCR-like binding modes are enforced by fine-tuning AF2 on TCR:pMHC structures and binding data (*14, 15*) and by providing AF2 with generic TCR-like docking templates as input features (see Methods). The re-docking step allows the binding mode to adjust to the designed CDR sequences and provides AF2 quality metrics for design ranking. To obtain a second view of design quality, a version of the RoseTTAFold2 (RF2) network (*16, 17*) fine-tuned on antibodies and TCRs is used to model the final design sequence, providing a confidence estimate as well as a predicted structure that can be compared with the original design. This process is repeated many thousands of times with different template framework structures and different randomly selected CDR3 pairs, resulting in a large ensemble of candidate designs that are ranked by a quality score that combines the AF2 and RF2 confidence scores as well as the similarity between their structural models (see Methods). The top-ranked 100-500 designs for each pMHC target are then taken as starting points for a genetic algorithm (GA) refinement procedure, conducted in parallel across multiple concurrent processes operating on a shared design pool (**Fig. 1**, bottom). Each process repeatedly selects a random design from the current pool, mutates two interface positions, redocks and redesigns the interface, and then replaces an existing design in the current pool if the refined design has superior metrics (with limits to prevent a single lineage from taking over the pool, see Methods and **Figure S1**). ADAPT refinement can be viewed as iterative alternation between ProteinMPNN sequence design and AF2 redocking, with interface mutations introducing diversity and the GA framework enabling intensified exploration of promising regions of design space.

**Figure 1.**
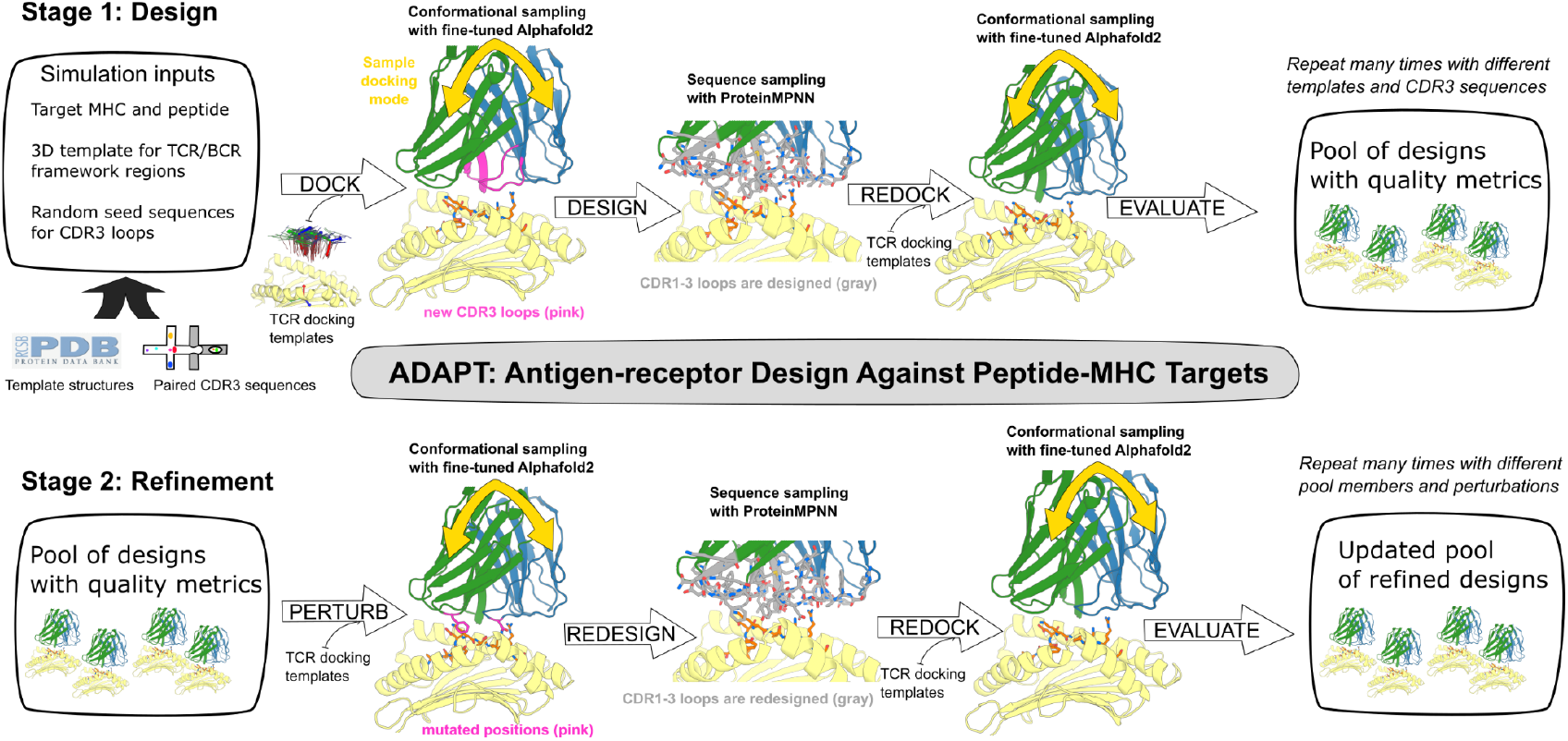
The ADAPT pipeline. In the design stage (top), many independent design simulations are conducted with different template structures for the TCR or antibody framework regions and different starting CDR3 sequences. Together with the pMHC target, these inputs determine the complete sequence of the system, which is modeled with a version of Alphafold2 (AF2) that was fine-tuned on TCR:pMHC structures and binding data (“Dock” arrow). Sampling of canonical docking geometries is encouraged by providing AF2 with four generic TCR:pMHC docking orientations spanning the observed docking modes in native structures (“TCR docking templates”). After docking, the sequence of the CDR1, 2, and 3 loops is optimized with the ProteinMPNN algorithm (“Design”). The structural model is then updated (“Redock”), and design quality is assessed using AF2 and RoseTTAFold2 (“Evaluate”). In the refinement stage (bottom), the highest quality designs are iteratively refined by a genetic algorithm that includes random interface mutations followed by AF2 redocking (“Perturb”) and CDR sequence redesign (“Redesign”).

We applied this TCR design pipeline to a panel of 9 pMHC targets including cancer-associated epitopes, tumor neoepitopes, and viral peptides (**Table 1**). Top-ranked sequence designs were introduced into TCR-null Jurkat T cells and assessed for binding to on- and off-target pMHCs by flow cytometric analysis with multimerized pMHC staining reagents (see Methods). For four targets (A01-EVD, A02-LLW, A02-GLM, and A02-TLM; **Table 1**), successful binder designs were identified in small-scale clonal screens consisting of 30-60 sequences.

**Table 1.**
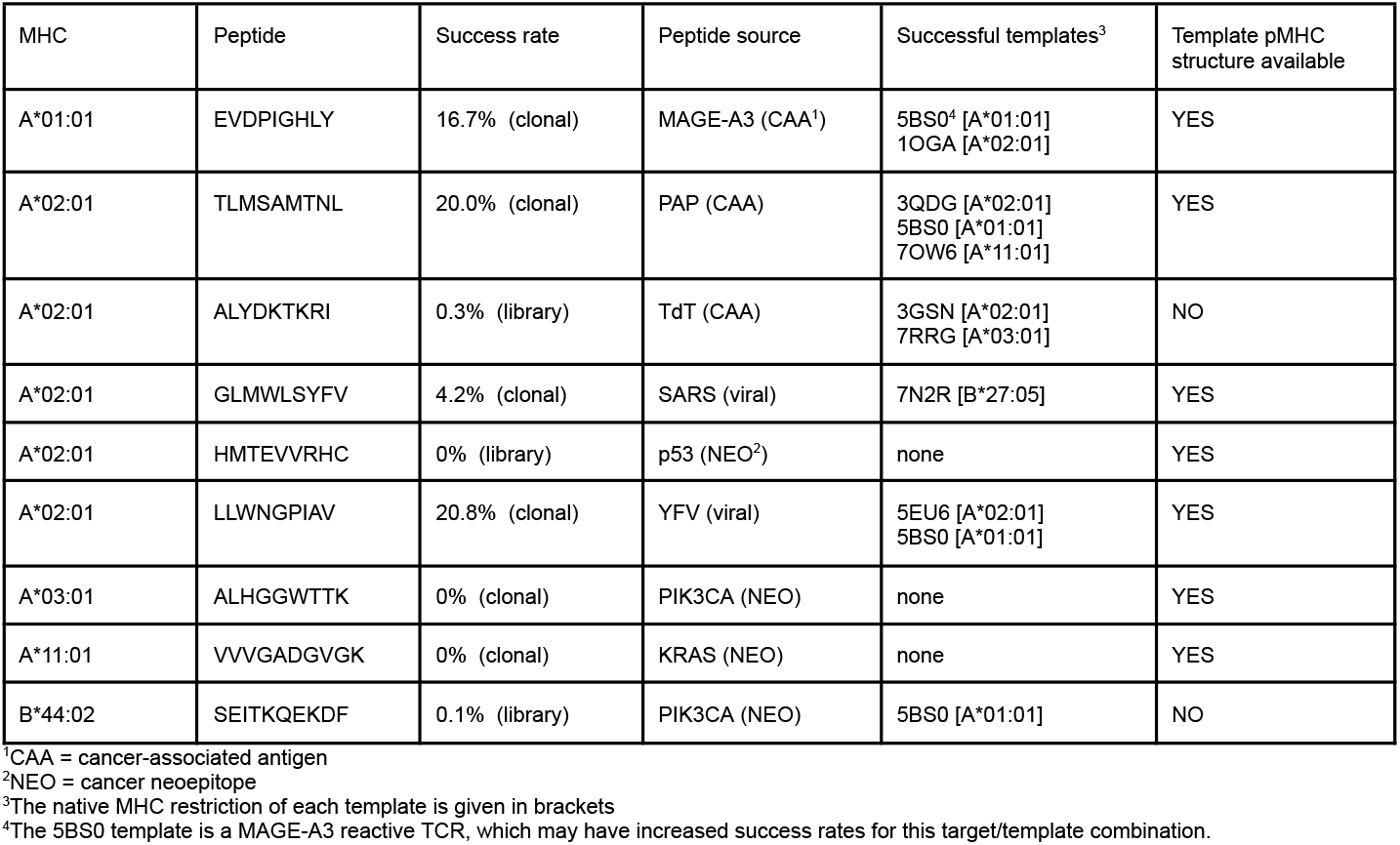
TCR design targets and outcomes.

Binders for two additional targets (A02-ALY and B44-SEI) were found by screening larger libraries of 1000-5000 designs built by cloning chip-synthesized DNA fragments into template-specific backbones (see Methods). No successful designs were identified for the remaining three targets (A02-HMT, A03-ALH, and A11-VVV). Structural models and binding data are shown in **Figure 2A** for a single design for each of the six successful targets. On- and off-target binding for 35 nonredundant designs from 17 distinct refinement lineages, along with a dendrogram showing their sequence relationships, are summarized in **Figure 2B** and **Figure S2**. The scatter plot in **Figure 2C** shows binding data for a larger set of successful and failed designs; note that the off-target pMHCs span a range of similarities to the design target, from the highly similar Titin-derived off-target for A01-EVD to more distant and therefore less challenging discrimination targets (**Table S1**). The designed CDR sequences are highly mutated relative to their template structures, with 70-80% of the designable positions changing in a typical loop (**Table S2**). To visualize the diversity of binding modes in these designs, we calculated the 6 rigid-body orientational parameters relating pMHC and TCR for each design and for a non-redundant set of MHC class-I restricted native TCR structures (see Methods). Visualization of the docking geometries by principal components analysis (PCA) shows that the designs are well distributed across the landscape of native TCR binding modes (**Fig. 2D**) and that designs sample binding modes that are distinct from those of the experimental structures used to template their framework regions (**Fig. S3**). Retrospective modeling of the designed sequences with the default Alphafold3 (*18*) pipeline (without interface templates or fine-tuning) recapitulated the designed binding geometries with high confidence (**Fig. S4A**). Finally, we performed an all-vs-all binding screen matching the six representative designs against all six target pMHCs and observed binding to the intended target along with some off-target binding, particularly to the A02-TLM pMHC (**Fig. 2E**). Alphafold3 metrics showed a substantial correlation with experimental binding results (**Fig**. **S4B-C, Fig. S5**). Taken together, these results demonstrate that the ADAPT pipeline can generate diverse, novel TCRs that bind to their intended pMHC targets.

**Figure 2.**
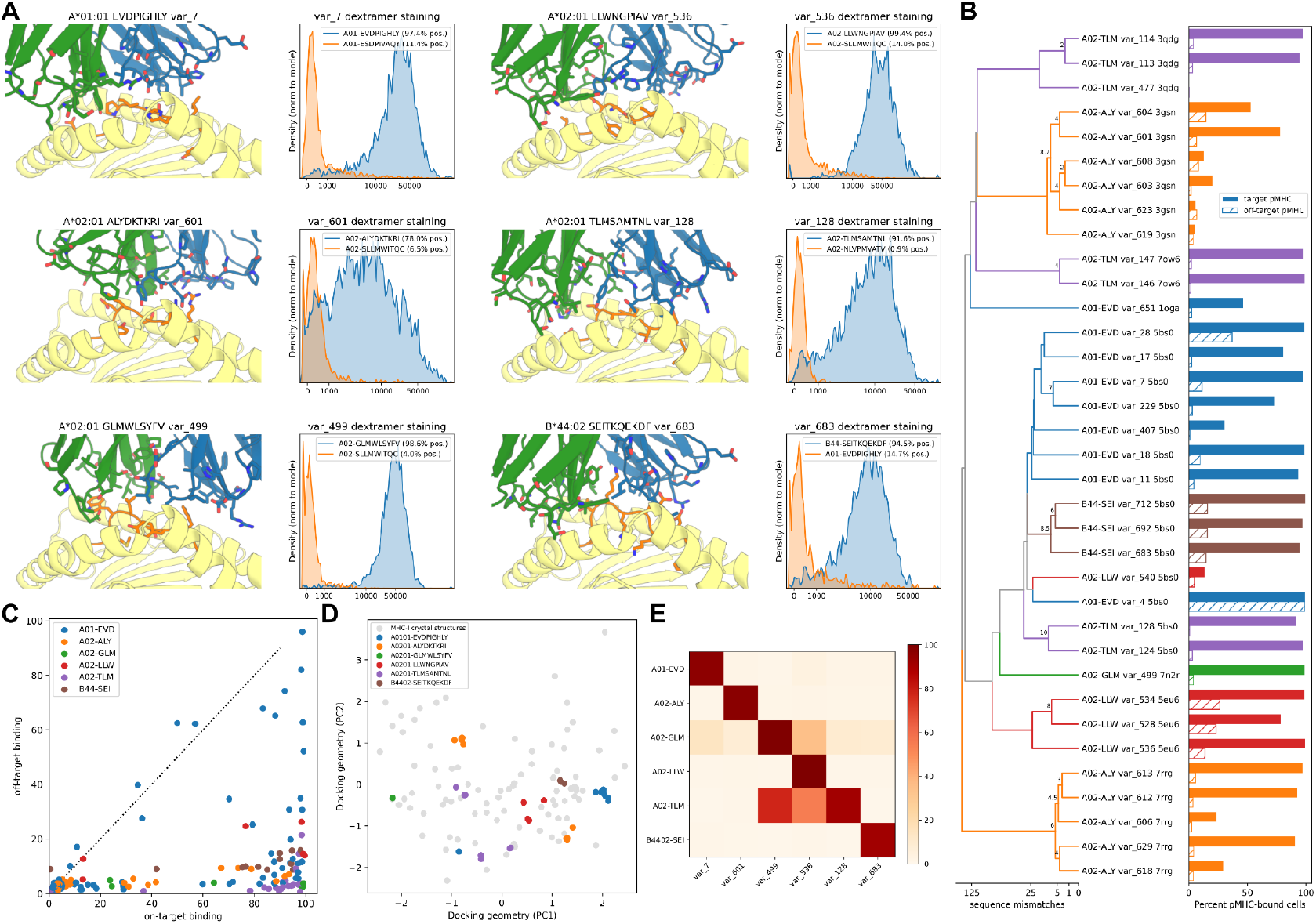
Designed TCRs bind their targets. **(A)** For six representative designed TCRs, one from each successful pMHC target, are shown design models (MHC in yellow, peptide in orange, TCRα in green, and TCRβ in blue) and flow cytometry binding histograms for on-target (blue) and off-target (orange) pMHC multimers. **(B)** Summary bar plots of on- and off-target binding for 35 selected designs along with a dendrogram of their sequence relationships. **(C)** Scatter plot of on- vs off-target binding for a larger set of successful and failed designs, colored by pMHC target. **(D)** Docking geometry landscape of native TCR:pMHC interfaces (gray points) and designed TCR interfaces (colored by pMHC target) visualized by projecting the 6-dimensional rigid-body transform space to its top 2 principal components (PC1 and PC2). **(E)** Heatmap showing all-vs-all flow-cytometric binding data (percent cells bound by pMHC target) when the six representative designs are paired with all six pMHC targets.

Like TCRs, antibodies are generated through a genetic recombination process that produces diverse CDR loop regions which mediate target binding. Because they function in part as secreted proteins, antibodies often make better soluble reagents than TCRs, which remain anchored to the T cell surface. Thus pMHC binding antibodies might have advantages over TCRs when a soluble targeting molecule is desired, for example as a component of a bispecific T cell engager (*19, 20*). To date, a handful of pMHC-binding antibodies have been identified by traditional immunization and screening approaches (*21–23*). By replacing the TCR template structures with a set of antibody variable region structures, ADAPT can design antibodies that target pMHC complexes using TCR-like binding modes. We used ADAPT to generate TCR-mimic antibodies that specifically recognize three different pMHC targets with relatively high success rates in small-scale screens (**Fig. 3A-C, Fig. S6, Table 2**). Monovalent affinities of these designs for their targets ranged from 5nM to 700 nM, with most designs binding in the high nanomolar range (**Table S3, Fig. S7**). Cryogenic electron microscopy (cryoEM) structures were determined for two of the designed complexes—vAB-30 bound to the A01-EVD pMHC and vAB-66 bound to A02-TLM—revealing atomic accuracy at the interface (**Fig. 3D-E**). C_α_-RMSD calculations over full complexes yielded values of 0.7Å and 0.6Å for vAB-30 and vAB-66 respectively; C_α_-RMSDs of 0.6Å (vAB-30) and 1.2Å (vAB-66) were calculated for the CDR loop regions after aligning on the MHC structures. For the vAB-30 complex, peptide recognition is primarily mediated by the heavy chain CDR2, with a hydrogen bond between Arg54^HC^ and the C-terminal Tyr9 main-chain oxygen. Additional stabilization arises from a Pro–aromatic CH–π interaction between Light chain CDR1 Tyr32^LC^ to peptide residue Pro4.

**Table 2.** Antibody design targets and outcomes.

| MHC | peptide | Success rate | Successful templates<sup>1</sup> | Successful binding orientations | RMSDs to cryoEM structure<sup>2</sup> |
| --- | --- | --- | --- | --- | --- |
| A*01:01 | EVDPIGHLY | 6.25% (clonal) | 6YIO [CD25]<br>7LSG [TBEV EDIII] | Forward | vAB-30: 0.7Å, 0.6Å |
| A*02:01 | TLMSAMTNL | 8.33% (clonal) | 7YV1 [K-Ras] | Reverse | vAB-66: 0.6Å, 1.2Å |
| A*02:01 | ALYDKTKRI | 5.21% (clonal) | 7KQL [Tim-3]<br>7SSC [HCMV peptide]<br>7YV1 [K-Ras] | Forward + Reverse | - |

<sup>1</sup>The target of the wild-type antibody template is given in brackets. None of the wild-type targets are pMHC complexes.
<sup>2</sup>The global Ca-RMSD and the Ca-RMSD for the CDR loops after aligning on the MHC

**Figure 3.**
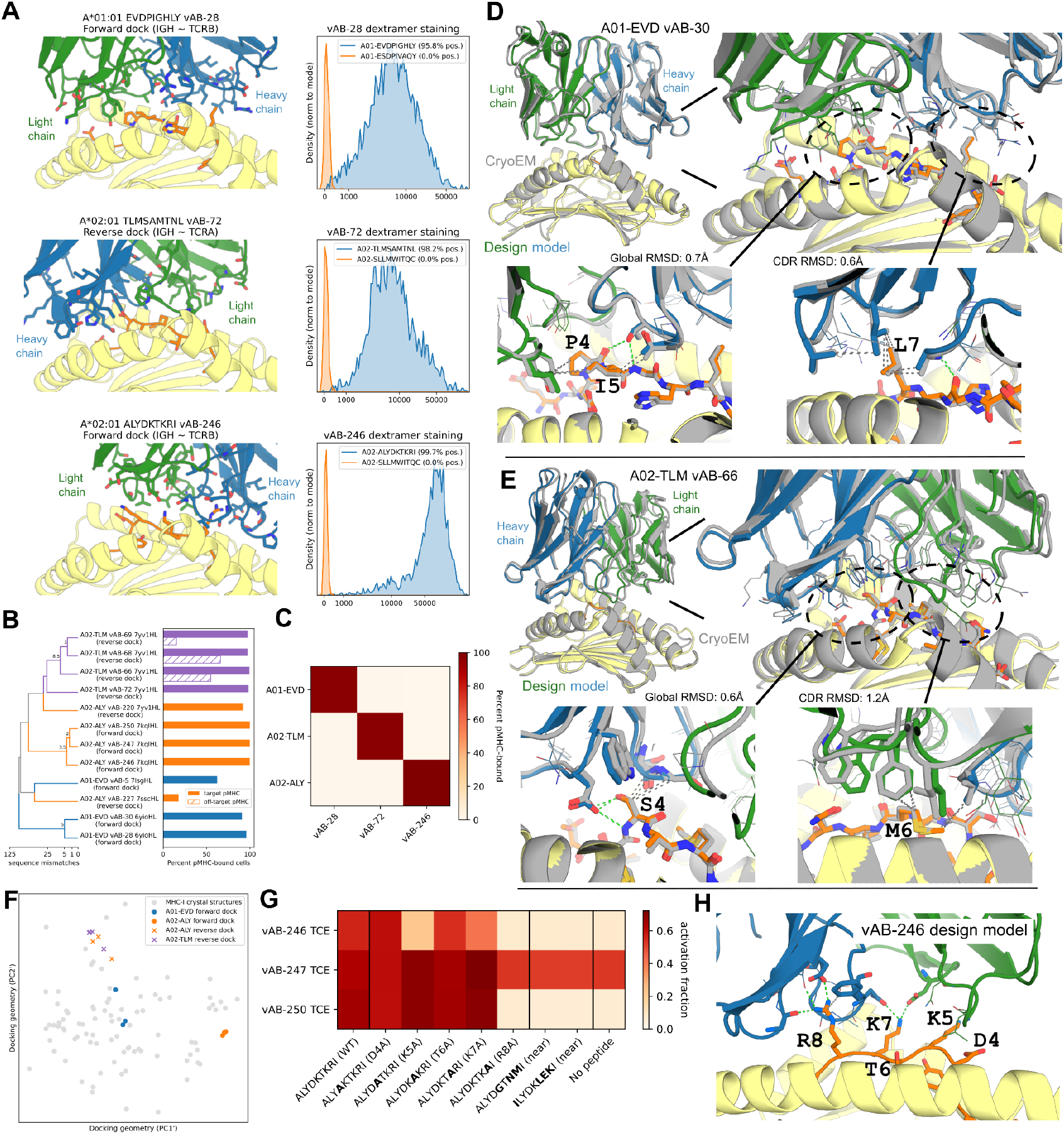
Designed antibodies bind their targets. **(A)** For three representative designed antibodies, one from each pMHC target attempted, are shown design models and flow cytometry binding histograms for on-target (blue) and off-target (orange) pMHC multimers. **(B)** Summary bar plots of on- and off-target binding for 13 selected designs along with a dendrogram of their sequence relationships. **(C)** Heatmap showing all-vs-all flow-cytometric binding data (percent cells bound by pMHC target) when the three representative antibodies are paired with all three pMHC targets. **(D)** Structural comparison of cryoEM structure (in gray) and design model (MHC in yellow, peptide in orange, heavy chain in blue, and light chain in green) for designed antibody vAB-30 bound to the HLA-A*01:01-EVDPIGHLY pMHC complex. **(E)** Structural comparison of cryoEM structure and design model (same colors as in panel D) for designed antibody vAB-66 bound to the HLA-A*02:01-TLMSAMTNL pMHC complex. **(F)** Docking geometry landscape of native TCR:pMHC interfaces (gray points) and designed antibody:pMHC interfaces (colored by pMHC target) visualized by projecting the 6-dimensional rigid-body transform space to its top 2 principal components (PC1’ and PC2’) after excluding rotations about the antibody pseudosymmetry axis. The excluded rigid-body dimension differentiates between forward-docking antibodies (disks) and reverse-docking antibodies (x’s); canonical native TCR:pMHC complexes are forward docking). **(G)** Heatmap of induced activation for bi-specific T cell engager (BiTE) constructs incorporating three designed antibodies (rows) in the presence of presenter cells pulsed with various target peptide variants (columns). **(H)** Design model for vAB-246 antibody bound to A02-ALY pMHC shows more contacts at peptide positions whose mutation to alanine reduces T cell activation in the presence of the vAB-246 BiTE construct (R8 and K7 versus T6 and D4).

Contacts to the MHC are distributed, involving the α1 helix from CDR2^HC^, CDR3^HC^ and α2 helix from CDR1^LC^, CDR2^LC^, and CDR3^LC^ **(Fig. S8A**). For the vAB-66 complex, peptide recognition is driven by the heavy chain CDR1, with a hydrogen bond from Asp33^HC^ to peptide residue Ser4, as well as van der Waals contacts from CDR3^HC^. Additional contacts from Phe32^LC^ and Trp92^LC^ extend the reach toward the C terminus of the peptide. Contacts to the MHC are more distributed, with contact to the α1 helix from CDR3^HC^, CDR1^LC^, and CDR2^LC^, and contact to the α2 helix from CDR1^HC^ (**Fig. S8B**).

Existing structures of antibodies with TCR-like recognition modes have shown two broad classes of docking geometry which are related by 180° rotation of the antibody about its pseudo-symmetry axis: one in which the antibody heavy chain aligns with the TCR beta chain (‘forward’ docking), and one in which the heavy chain aligns with the TCR alpha chain (‘reverse’ docking). This is controlled in the ADAPT pipeline at the docking and redocking steps (**Fig. 1**) through the use of docking-geometry templates that are aligned to the template antibody structure in either the forward or reverse orientation. The forward or reverse orientation is randomly selected at the start of each individual design calculation in order to sample candidate designs with both orientations. As shown in **Figure 3** and **Table 2**, the successful pMHC binders exhibit both orientations. To visualize the designed binding modes relative to native TCRs, we performed PCA analysis of their orientational parameters, now dropping the single parameter describing rotation of the antibody about its internal pseudosymmetry axis (which corresponds to the choice of forward vs reverse dock; see Methods). This analysis showed that the designed binding modes fall within the space sampled by natural TCRs (**Fig. 3F**).

To explore potential clinical applications, we formulated three of the A02-ALY specific antibodies as bi-specific T cell engagers (see Methods) and evaluated their ability to trigger T cell activation in the presence of target cells pulsed with peptide variants. Two of the designs (vAB-246 and vAB-250) showed specificity for the target peptide and its near variants (**Fig. 3G**), with the pattern of sensitivity to peptide mutations for vAB-246 appearing to correlate with extent of antibody contact at the designed interface (**Fig. 3H**). The third design induced activation even in the absence of pulsed peptide, suggesting that it binds to off-target pMHC complexes on the target cell surface.

For application as cytotoxic targeting reagents, TCRs must have high specificity toward their target epitope in order to avoid killing healthy cells bearing structurally similar self-pMHCs. This is challenging because the TCR recognizes a composite interface consisting of a few residues from the target peptide together with multiple MHC residues that are shared with many self-pMHCs. T cell activation provides a sensitive read-out of TCR specificity due to the signal-amplifying properties of the activation signaling cascade (*24*). Using transgenic T cells expressing designed TCRs co-cultured with antigen presenting cells in the presence and absence of pulsed target peptide, we found that many of the designed TCRs—despite being able to discriminate their target peptide from specific off-target competitors in binding screens—showed relatively high levels of non-specific activation in the absence of target peptide (**Fig. 4A-B; Fig. S9**); the more specific designs tend to have weaker on-target binding than non-specific designs (**Fig. 4C**).

**Figure 4.**
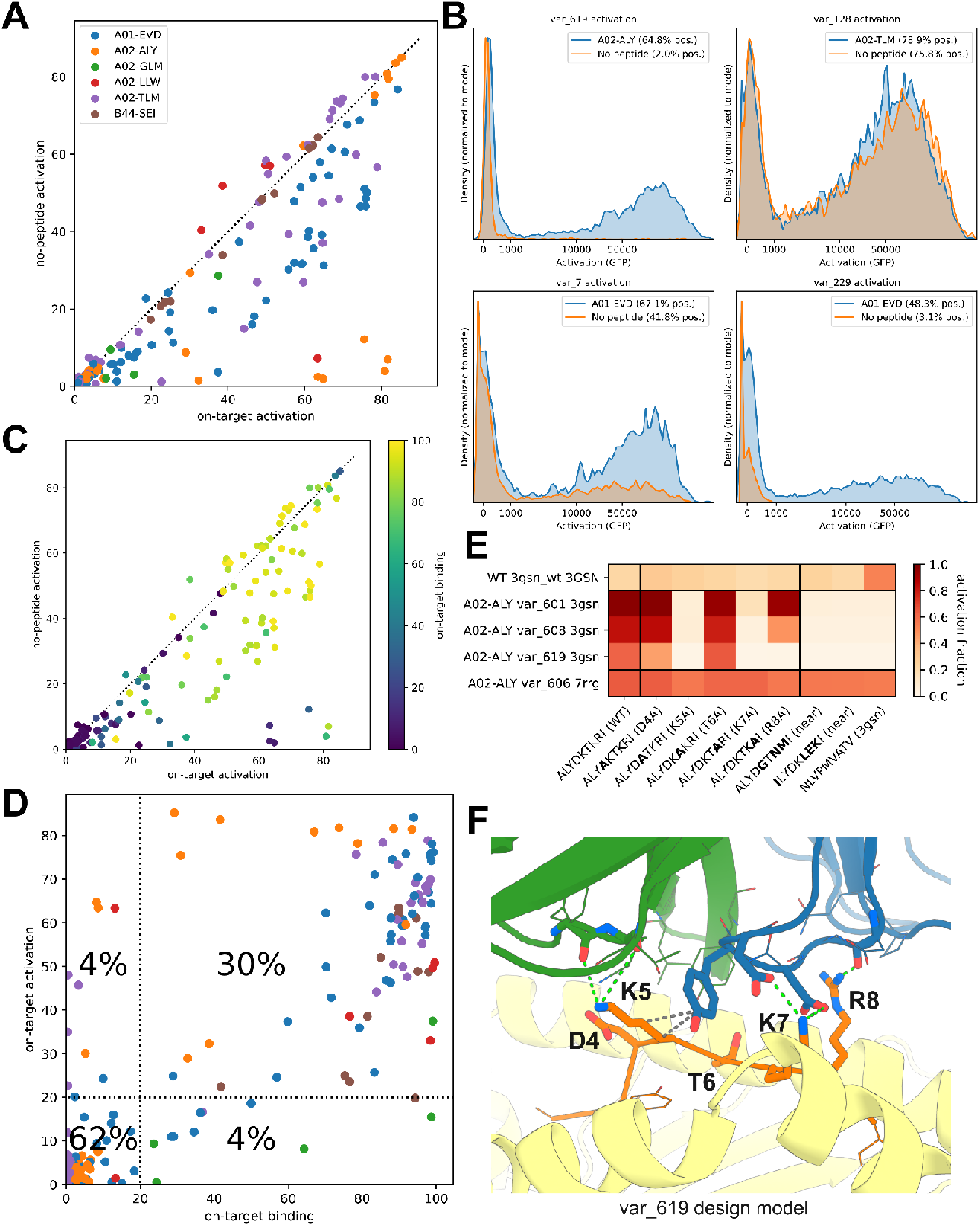
Designed TCRs activate T cells. **(A)** Scatter plot (colored by pMHC target) of T cell activation (percentage of transgenic T cells expressing a GFP activation marker) measured by flow cytometry for T cell populations expressing individual TCR designs in the presence of presenter cells pulsed with target peptide (x-axis) or presenter cells alone (y-axis). **(B)** Flow cytometry cell density histograms of activation marker expression for four designs (named in panel titles) with varying levels of off-target activation in the absence of target peptide. **(C)** Scatter plot of T cell activation in the presence or absence of pulsed target peptide colored by a measure of on-target binding strength for the corresponding TCR design (percentage TCR-transgenic cells bound by multimerized pMHC target). **(D)** Scatter plot of on-target binding (percentage multimer-bound cells) versus on-target activation (percentage activation-marker expressing cells) shows positive correlation across different TCR designs (colored by pMHC target). **(E)** T cell activation percentage for different A02-ALY-targeting designed TCRs or a wildtype antiviral TCR (rows) as a function of pulsed peptide identity (columns). Pulsed peptides include (left to right) the ALY target peptide from TdT, 5 alanine variants at positions 4-8, similar peptides from the human proteome, and the viral peptide target of the TCR template used for several of the designs (PDBID 3gsn). Rows labeled with the target, design name, and TCR template on which the design was based (3gsn or 7rrg) or “3gsn_wt” for the wildtype antiviral TCR from the 3gsn PDB structure. **(F)** The design model of the var_619:pMHC interface shows that peptide positions where alanine mutations disrupt activation (K5/K7/R8) tend to have more interface contacts than positions like T6 where mutations are tolerated.

Overall, there was a strong positive correlation between on-target binding and on-target activation: designs that bound above a threshold tended to support T cell activation and vice versa (**Fig. 4D;** Pearson’s *r* = 0.87).

Several designs targeting the highly polar HLA-A*02:01-restricted epitope ALYDKTKRI from terminal deoxynucleotidyl transferase (TdT) showed low levels of non-specific activation (lower right corner of **Fig. 4A**). We evaluated their fine specificity by additional activation experiments with peptides containing individual alanine mutations and similar peptides from the human proteome (**Fig. 4E**). Several designs were able to discriminate the target peptide from some, though not all, of the nearby mutant peptides, and the pattern of mutation sensitivity could be rationalized by examination of the designed structures, with mutations at non-contacted peptide positions being better tolerated than mutations at positions with extensive TCR contacts (**Fig. 4F**).

## Discussion

The specificity of T cell receptors for their pMHC epitopes underpins the exquisite precision of the adaptive immune system, providing protection from pathogens and elimination of transformed cells. Computationally designed TCRs and antibodies that specifically recognize tumor-associated pMHCs could form the basis of soluble or cell-based cancer immunotherapies; designs targeting pathogen-derived epitopes or immunogenic self-peptides could have applications treating infectious or autoimmune diseases, respectively. There has been exciting recent progress designing mini-protein binders for pMHC targets (*25*–*27*), yet there remain compelling reasons to design TCRs and antibodies for this purpose: engineered TCRs introduced into transgenic T cells benefit from fully native signaling pathways; there are well-established pipelines for development of antibody therapeutics; TCRs and antibodies may be less immunogenic than fully synthetic mini-proteins; and TCR design may yield insights into the function of natural TCRs, with implications for disease diagnosis and basic immunology. Our structure-based pipeline for the computational design of TCRs and antibodies that bind to pMHC targets iterates between conformational sampling with a fine-tuned structure prediction network and fixed-backbone sequence design. Application of this approach to nine distinct pMHC targets produced TCR binders for 6 of them, 4 from small-scale clonal screens. Two of the successful targets lacked a previously determined pMHC structure (**Table 1**), which suggests that it should be feasible to target new and less well-characterized epitopes. A simple extension to antibody design yielded binders from small screens for all three targets attempted, including a highly polar peptide (ALYDKTKRI) in complex with HLA-A*02:01. The pipeline’s success in designing challenging, loop-mediated interactions at relatively high success rates demonstrates that structure prediction algorithms like AF2 can be powerful sampling engines for protein design, provided that sampling can be focused on relevant conformations (here through the use of fine-tuning and pMHC docking templates) and that sufficient diversity can be introduced (in our case through the use of diverse seed CDR3 sequences taken from natural human TCRs). Requiring only two forward passes through the AF2 network per design or refinement simulation (**Fig. 1**), the ADAPT pipeline has modest GPU hardware requirements relative to more memory-intensive gradient-backpropagation approaches (the designs reported here were generated on commodity NVIDIA GPUs with 11 GB of VRAM). We additionally found in preliminary testing that ADAPT was better able to generate native-like TCR docking modes than a fine-tuned version of RFdiffusion (**Fig. S10**), which may suggest that structure-prediction based design approaches can be advantageous in geometrically-constrained settings with multiple, diverse examples for training.

The biophysical determinants of T cell activation are incompletely understood (*28*). Our large panel of ADAPT-designed TCRs, generated without the influence of positive or negative thymic selection, provides insight into the relationship between TCR binding and T cell activation. We found a significant (*p*<10^-80^) positive correlation (**Fig. 4D**) between the levels of on-target binding, as measured by staining with multimerized pMHC reagents, and on-target activation from peptide-pulsing experiments. With some exceptions, designs that showed measurable binding to their targets also mediated T cell activation by presenter cells pulsed with target peptide. Given that all of these TCRs were designed to bind in canonical docking modes within the envelope of previously characterized TCR structures (**Fig. 2D**), our results show that binding with a canonical docking mode can support TCR activation (*29, 30*). Within the range of affinities sampled by these designs, we do not see strong evidence for additional biophysical or interface chemistry constraints beyond canonical binding.

While the ability of our design pipeline to generate TCRs that bind pMHCs and activate T cells is an advance for the field, there is still considerable room for improvement to generate more specific designs. The specificity of nearly all of these in silico-generated sequences is likely insufficient for targeted killing of specific cell populations, and hence further optimization by experimental or computational means will be necessary. Our pipeline could likely be improved by favoring contacts to the peptide over the MHC and by encouraging more native-like sequences in the CDR1 and CDR2 loops (which are currently being enriched for hydrophobic amino acids, **Fig. S11**). More broadly, our difficulty in designing TCRs that activate exclusively in the presence of a target peptide highlights the importance of negative thymic and peripheral selection steps in the development of natural T cells.

## Methods

### Computational design methods

In the first stage of the pipeline, many independent docking and design calculations are carried out, each with a different random choice of template antigen-receptor (TCR or antibody) structure and initial CDR3 loop sequences (**Fig. 1**, top). The template receptor structure determines the sequence and structure of the non-CDR framework regions and of the CDR1 and CDR2 loops. TCR template structures were taken from the RCSB protein databank (*31*). Antibody template structures and metadata were downloaded from the SAbDab database (*32*). The CDR3 loop sequences, taken from a paired human T cell receptor, are spliced into the template sequence to create a hybrid receptor sequence that is provided to the fine-tuned AF2 network along with the sequences of the target MHC and peptide. AF2 is run without MSA input but is provided with four multichain templates assembled by combining sequence-similar peptide-MHC structures together with the structure of the template receptor positioned in four different docking modes relative to the pMHC. These four ‘generic’ docking geometries, which are fixed and identical for every design calculation, were chosen to minimize the rigid-body distance to a set of canonical ternary TCR:pMHC structures from the RCSB protein databank (*31*). In other words, the four generic docks were chosen to optimally cover the space of potential docks as inferred from experimentally determined TCR:pMHC structures. The database of paired CDR3 sequences contains 1,688,863 sequences assembled from publicly available single-cell genomics studies on T cells (*33*). The fine-tuned AF2 model was trained on experimentally-determined TCR:pMHC structures and binding data using the “alphafold_finetune” approach described in Ref. (*15*) (https://github.com/phbradley/alphafold_finetune).

After AF2 docking, the sequences of the 6 CDR loops (CDRs 1-3 on both the alpha/light and beta/heavy chains; IMGT (*34*) loop definitions are used) are redesigned using the ProteinMPNN network (*12*) with the following parameters: temperature=0.1, model_name=‘v_48_020’, num_seq_per_target=3, number of edges=48, training noise level=0.2Å. After sequence design, a new docked structure is generated using AF2 with docking templates exactly as in the initial docking step. This structure represents the final design model, and the AF2 quality metrics are recorded for selecting the top designs.

To evaluate design model quality, the mean AF2 predicted aligned error (PAE) score for all (pMHC,TCR) and (TCR,pMHC) residue pairs is computed. The designed sequence is also input to a RoseTTAFold2 (RF2) model fine-tuned on antibody and TCR complex structures (*16*), and the CDR3 RMSD between the design model and the RF2 prediction is calculated after aligning on the MHC chains. Scatter plots of these design quality metrics for representative design runs are shown in **Figure S1**. A single weighted ranking score is calculated according to the formula:

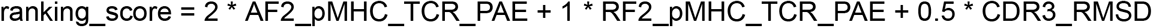

Minimizing this score selects for lower TCR-pMHC PAE values (ie, higher structural confidence) and lower RMSDs (ie, greater similarity) between the AF2 and RF2 models of the designed interface. The CDR3 RMSD value is calculated after aligning both models on the MHC chain, so it captures both the internal conformation of the CDR3 loops as well as their docked orientation relative to the MHC.

In the second stage of the pipeline, the top 100-500 designs by ranking score are selected to form an initial pool for subsequent refinement with a parallel genetic algorithm. During the refinement process, multiple independent simulations operate independently on the pool of designs, each selecting targets for refinement from the current pool at random and adding the refined models to the pool if they improve it, subject to a diversity constraint that prevents a single lineage from taking over the pool. This is necessary because refined designs do not replace their parent in the pool, which allows for intensified sampling in promising regions. The default diversity constraint caps the number of designs allowed in the pool from the same lineage at 10. Here a single lineage is defined as the set of all designs descended from the same founding member of the refinement pool.

Each independent refinement trajectory consists of a perturbation, a redesign, and a redocking calculation. Perturbation consists of randomly mutating two of the CDR positions to new amino acids and redocking with AF2. All positions in the six CDR loops are then redesigned with ProteinMPNN, and a final model with quality metrics is generated with AF2 redocking and RF2 reprediction. At this point the current refinement pool is read back into memory and any refined designs that are superior to existing pool members replace those old designs (subject to the lineage diversity constraint). Thus a fixed pool size is maintained during the refinement process.

### Docking Geometry Analysis

The TCRdock software (*14*) (https://github.com/phbradley/TCRdock) was used to assign six rigid-body docking geometry parameters to each design model and to a set of nonredundant native class I TCR:pMHC ternary structures. In this analysis, reference coordinate frames are first defined for the MHC and for the TCR. In both cases, the reference frame is defined by a set of residue pairs related by an approximate C2 symmetry axis. In the case of the MHC, these residues are taken from the antiparallel beta sheet that forms the floor of the peptide binding groove (**Fig. S12A**). For the TCR, they consist of a set of 13 structurally conserved framework positions that superimpose closely when aligning the TCR alpha and beta chains (or their analogous aligned positions in the antibody light and heavy chains) (**Fig. S12B-C**). In each case, the x-axis of the reference frame is defined by the axis about which a rotation optimally superimposes the residue pairs (is, the approximate symmetry axis of the residue pairs), with the frame origin and z-axis defined by the centers of mass of the residue sets being aligned (**Fig. S12D**). With these choices, the x-axes of the TCR and MHC frames are approximately aligned and antiparallel (red arrows in **Fig. S12D**). Once the frames are defined, the six orientational parameters are taken to be (1-2) the y- and z-coordinates of a unit vector from MHC to TCR represented in the MHC local coordinate system, (3-4) the y- and z-coordinates of a unit vector from TCR to MHC represented in the TCR local coordinate system, (5) the distance between the reference frame origins, and (6) the dihedral angle formed by the y-axis vectors along the rotation axis connecting the frame origins.

For TCRs binding in a canonical orientation, this last parameter clusters around 180 degrees, whereas for TCR-mimic antibodies, dihedral angles of ∼0 and ∼180 degrees are observed, corresponding to the reverse and forward docking modes, respectively.

### Lentivirus preparation and transduction

Lentiviral particles were produced in HEK293T cells (ATCC, CRL-3216) using a standard third-generation packaging system. Cells were seeded to reach ∼80% confluence after 24 h and transfected with pMD2.G (Addgene #12259), psPAX2 (Addgene #12260), and the lentiviral transfer vector at a 2:3:4 plasmid mass ratio using Lipofectamine 3000 (Thermo Fisher Scientific, L3000015) in Opti-MEM. 800 and 150 ng total DNA was transfected for 24-well and 96-well scale. After 4 h, the transfection medium was replaced with fresh DMEM containing 10% FBS and 1x penicillin–streptomycin (Thermo Fisher Scientific, 15140122) or Antibiotic-Antimycotic (Thermo Fisher Scientific, 15240062). Viral supernatants were harvested 36–48 h post-transfection, clarified through 0.45 µm low-protein binding filters, used fresh or aliquoted, and stored at -80 °C.

For transduction, target cells in log growth phase were resuspended in complete medium optionally supplemented with 8 µg/mL polybrene (Santa Cruz Biotechnology, sc-134220) and mixed with 5 to 200 uL of fresh or thawed lentivirus. Optionally, cells were spinned at 1000 × g for 2 h at 33 °C, then resuspended and incubated under standard culture conditions. Media were refreshed after 3 h, and cells were passaged or expanded 24 h post-infection. Puromycin selection (Thermo Fisher Scientific, A1113803) was initiated 48 h after transduction at 500-1000 ng/mL. Transduction efficiency was assessed by flow cytometry for mTagBFP2 expression 48 h post-transduction.

### Transposon based stable cell line production

iOn plasmids (PMID: 32559415) were co-transfected with pCAG-hyPBase at a 3:1 molar ratio using the Lonza 4D-Nucleofector system (Lonza). For each reaction, 4 × 10^5^ cells were used per well in 20 µL nucleocuvettes with a total of 2 µg DNA and nucleofected according to the manufacturer’s recommended program for the respective cell type. Following nucleofection, cells were transferred to pre-warmed complete medium and incubated under standard culture conditions. Cells were expanded the next day and Puromycin selection started at 500 ng/mL 48 hours post nucleofection. For library nucleofection, a number of 8 reactions were performed for a total of 3.2 × 10^6^ nucleofected cells to achieve a coverage of at least 100x of the library complexity.

### Variant plasmid preparation

Plasmids containing TCR and BCR variants used in clonal assays were either obtained from Genscript, IDT, or Twist, or cloned in house by cloning DNA fragments containing the coding sequence into receiving vectors using golden gate assembly with BsaI. When cloned in-house, ZymoPure Plasmid Miniprep kit (Zymo Research) was used.

### Pooled plasmid cloning

The original iOn and lenti backbone were modified to enable compatibility with BsaI, BsmBI, PaqCI BbsI, and SapI. Due to the higher cloning efficiency, the iOn plasmid was used for assembling pooled libraries. Pooled libraries were obtained as linear fragments from Twist (Multiplexed Gene Fragment product). Each DNA fragment contained four designed sequence regions (CDR1b–FW2–CDR2b, CDR3b, CDR1a–FW2a–CDR2a, and CDR3a), arranged in order and separated by three pairs of antiparallel Type IIS restriction sites (BsmBI, PaqCI, and BbsI) to enable directional Golden Gate assembly. The entire construct was flanked by outward-facing BsaI sites and subpool-specific adapter sequences for PCR amplification and library integration (**Fig. S13**). After receiving the double stranded library, we performed qPCR to amplify the subpools corresponding to designs made using a given framework against a specific target. Amplified subpools were either gel extracted or purified using SPRIselect magnetic beads (Beckman Coulter) and quality controlled with gel electrophoresis. We then performed golden gate assembly, PCR clean-up (DNA Clean and Concentrator-5 by Zymo Research), electroporation into Endura electrocompetent *E. coli* (Biosearch Technologies) and grown overnight in 37°C in shake flasks containing media with antibiotics before harvesting and plasmid prep. We used Plasmidsaurus whole plasmid sequencing to quality control each prep.

### Cell based binding assay

Jurkat cells deficient in TRAC and TRBC containing a NFAT-RE_EGFP construct (Jurkat ab-) were transduced with lentivirus, undergone puromycin selection, and then stained with APC-conjugated anti-CD3 antibody (clone UCHT1 Biolegend) and PE-conjugated peptide MHC Dextramer (Immudex) or Tetramer (Fred Hutch immune monitoring core) and analyzed with an Attune NxT Flow Cytometer (Thermo Fisher).

TCR hits against B44-SEI targets were initially tested by transient transfection into HEK293T cells stably expressing CD3 ε, δ, ζ, γ, then stained and analyzed 48h post-transfection similar to Jurkat ab-cells.

### T cell co-culture stimulation assay

Jurkat ab-cells transduced with TCRs and selected for puromycin expression were harvested, washed, with fresh cRPMI and co-incubated with appropriate antigen presenting cells in a ratio of 1:1 in the presence of anti-CD49d (BD Biosciences) and anti-CD28 (BD Biosciences) each at a 1:1000 dilution. Then, appropriate peptides were pulsed in at the desired concentration (with 1 uM as the most common concentration used) and incubated overnight at 37°C with 5% CO_2_. Then the media was replaced with DPBS and the cells were analyzed with flow cytometry to determine the fraction of EGFP^+^ cells among mTagBFP2^+^ population. In experiments containing T cell engagers (TCEs) the appropriate TCE protein was added at the desired concentration (50 nM as the most commonly used concentration).

### Screening scFvs through yeast display

We performed yeast surface display for screening some of the designed antibodies with the standard protocol described in Cao et. al. (*35*). The scFvs were fused through the N-term to Aga2p with a VH - GS Linker - VL orientation. Staining was performed using pMHC Dextramers or Tetramers and anti-Myc or anti-IgG1 antibodies.

### Mammalian display of antibodies

We created vectors for mammalian display of antibodies by fusing them to PDGFR transmembrane domain in either the iOn or Lenti backbone.IgG1-Fabs and full-length IgG1s were fused to PDGFR through the C-term of their heavy chain while the light chain was not fused to PDGFR. Staining was performed similar to staining Jurkats harboring TCRs.

### Bi-specific T cell engager production

Chimeric IgG based T-cell engagers were cloned by fusing an anti-CD3 scFv to the C-terminus of the human IgG1κ gene and swapping the canonical Fc domain with a silenced Fc variant (*36*). These variants were first screened for binding on the HEK293T surface using a displayed protein A based capture system. Variants with correct binding were moved forward and protein preps for these constructs were obtained from Genscript through CHO-S or Genewiz using Expi293 cell expression and Protein A based purification.

### Affinity measurement using Surface Plasmon Resonance (SPR)

Binding affinity measurements were performed on a Biacore 8K instrument (Cytiva) at 25 °C using HBS-EP+ buffer (10 mM HEPES pH 7.4, 150 mM NaCl, 3 mM EDTA, 0.05% v/v Tween-20). For the A01-EVD system, human IgG1s were immobilized on a Protein A sensor chip and serial dilutions of biotinylated peptide–MHC complexes (pMHCs) were injected as analytes. For all other targets, biotinylated pMHCs were immobilized on a streptavidin (SA) sensor chip and monovalent IgG1-Fab analytes were injected at multiple concentrations. Association and dissociation phases were monitored under continuous buffer flow, and reference subtraction (blank and control surfaces) was applied to remove nonspecific signal. Measurements were performed using single-cycle kinetics (sequential analyte injections without surface regeneration). Data were globally fitted using a 1:1 Langmuir binding model to extract kinetic rate constants (k_º_□, k_º_ff) and equilibrium dissociation constants (K_D). Monovalent Fabs for SPR were prepared by digestion of full-length IgG1s (acquired from Genscript or Genewiz) with LysC or Papain and removing the undigested or Fc portion with Protein A based agarose columns.

### Cryo-EM sample preparation and data collection

Fab proteins of vAB30 and vAB66 for structural characterization were obtained from Genscript. The designed variable domains were fused to the human CH1 domain of IgG1 including the hinge domain, cloned in pcDNA3.4, expressed in CHO-S cells and purified with CH1 resin.

For A01-EVD, the pMHC protein used for cryo-EM was expressed in Expi293 cells (Thermo Fisher Scientific). A single-chain trimer (SCT) construct (*37*) was cloned into the pD649 vector, consisting of the peptide, a (G_3_S)_3_ linker, β2-microglobulin, a (G_4_S)_4_ linker, the HLA-A*01:01 α chain, and a C-terminal His tag. A total of 200 µg of SCT plasmid was transfected into 400 million Expi293 cells following the manufacturer’s instructions. After four days, cells were pelleted at 500 × g for 10 min, and the supernatant was collected and diluted 1:1 with PBS. 4 mL Ni–NTA resin were added, and the mixture was incubated overnight at 4 °C with gentle rotation. The resin was collected on a gravity column, washed once with PBS (pH 7.2) containing 20 mM imidazole, and eluted with PBS (pH 7.2) containing 200 mM imidazole. The eluate was concentrated using a 30 kDa cutoff filter (Millipore) and further purified by size-exclusion chromatography on a Superdex 200 column (GE Healthcare) using an ÄKTA Purifier. Fractions containing the target protein were pooled for downstream analyses.

A02-TLM was refolded from inclusion bodies (IBs) by the as previously reported with modifications (*38*). HLA A*02:01 and β-2-microglobulin (β2m) were cloned into pET28a and pETDuet vectors and expressed separately in E. coli strain *BL21* as IBs. The cells were lysed by sonication in 50 mM Tris pH 8.0, 1% v/v Triton X-100 (Sigma Aldrich), 100 mM NaCl, 5 mM MgCl2, 10 mM DTT, and 0.2 mM PMSF, and benzonase. Following lysis, EDTA was added to 10 mM and IBs were pelleted by centrifugation at 10,000 g for 15 minutes. IBs were washed three times in buffer containing 50 mM Tris pH 8.0, 0.5% Triton X-100, 100 mM NaCl, 1 mM Na-EDTA, 1 mM DTT, and 0.2 mM PMSF, and once in the same buffer omitting the Triton X-100. IBs were solubilized in 8 M urea, 50 mM Tris pH 8.0, 0.5 M EDTA, and 1 mM DTT and stored as frozen aliquots.

To refold, 20 mg of TLM peptide (Genscript) in DMSO was added dropwise to 400 ml refolding buffer (100 mM Tris pH 8.0, 2 mM EDTA, 5 M urea, 500 mM L-arginine HCL, 0.2 mM PMSF, 1x protease inhibitor cocktail (Sigma)) stirring at 4°C. This was followed by addition of 0.5 mM oxidized and 5 mM reduced glutathione.

Aliquots containing 20 mg of HLA A*02:01 and of β2m were mixed and added dropwise. The refolding reaction was left stirring overnight, then dialyzed for two days at 4° C against 4 L of 100 mM Tris, pH 8.0, with three changes of buffer. The dialyzed protein was filtered with a 0.45 µm PMSF filter and a glass prefilter, concentrated using 10,000 MWCO centrifugal filters (Amicon), injected onto a Mono Q column (Cytiva) and eluted with a gradient to 400 mM NaCl. Protein-containing fractions were further purified on a Superdex S200 Increase column (Cytiva).

A plasmid encoding the β2m-targeting nanobody AD01 (*39*) was transiently transfected into expi293 cells. Supernatant was harvested after five days, passed over a Ni-IMAC resin (Thermo Scientific) and eluted with 250 mM imidazole in 150 mM NaCl, 10 mM HEPES pH 7.3 (HBS). Protein was concentrated and injected on a Superdex S75 Increase column (Cytiva) in HBS buffer.

Protein complexes for cryoEM were prepared by combining equimolar amounts of Fab, pMHC, and nanobody AD01, and purifying by size exclusion chromatography on a Superdex S200 Increase column (Cytiva). Complex formation and purity was assessed by SDS-PAGE.

To prepare cryo-EM specimens, 3.0 µL of each protein complex was applied to glow-discharged Quantifoil Au R1.2/1.3, 200-mesh (vAB-30) or 300-mesh (vAB-66) grids. Excess liquid was blotted for 1.0 s with filter paper, and grids were plunge-frozen in liquid ethane cooled by liquid nitrogen using a Vitrobot Mark IV (Thermo Fisher Scientific) operated at 8 °C and 100% humidity. Data were collected on a Titan Krios electron microscope (Thermo Fisher Scientific) operating at 300 kV and equipped with a Gatan K3 direct electron detector. Movies of the vAB-30/A01-EVD and vAB-66/A02-TLM complexes were recorded in super-resolution mode with SerialEM (*40*), at calibrated pixel sizes of 0.4135 Å and 0.94 Å, respectively. Patch motion correction was applied with binning to physical pixel sizes of 0.827 Å and 1.88 Å. Each movie stack contained 50 frames, with total exposures of 60 e^-^/Å^2^ (vAB-30) or 50 e^-^/Å^2^ (vAB-66), and defocus values ranged from -1.0 to -2.0 µm (vAB-30) or -0.8 to -2.0 µM (vAB-66). All movies were processed and quality-assessed in cryoSPARC Live.

For the vAB-30/A01-EVD dataset, 5,682 aligned micrographs were retained for further processing in cryoSPARC (*41*). A total of 4,636,242 particles were initially extracted using a 320/160 binned box. After two rounds of 2D classification, 1,864,082 particles were selected and subjected to non-uniform (NU) refinement, yielding a 3.4 Å reconstruction. Refinement employed multiple references, including one accurate and two biased models (*42*). Following duplicate removal, 1,317,109 particles remained. Re-extraction with a 320/240 binned box was followed by global CTF refinement, local CTF refinement, and NU-refinement, resulting in a final map at 2.6 Å resolution (**Fig. S14**). CryoEM maps have been deposited in the electron microscopy databank (EMDB) with accession EMD-73460 and movie stacks at the Electron Microscopy Public Image Archive (EMPIAR).

For the vAB-66/A02-TLM, 10,869 aligned micrographs were retained for further processing in cryoSPARC. An initial set of 639,144 blob-picked particles was 2D-classified and used as templates for picking a full 6,762,438 particle set extracted in a 256 pixel box downsampled to 72 pixels. This was reduced through two rounds of 2D classification to 1,807,378 particles, then by two rounds of heterogenous 3D refinement to 880,830 particles, with reextraction to a 256/200 pixel box. Particles were finally extracted without downsampling, followed by global CTF refinement, local CTF refinement, and NU refinement, resulting in a final map at 2.5 Å resolution (**Fig. S15**). CryoEM maps have been deposited in the EMDB with accession EMD-73458 and movie stacks at EMPIAR.

### Model building and refinement

For the vAB-30/A01-EVD structure, predicted atomic models of the complex were docked into the cryo-EM density maps, while for the vAB-66/A02-PAP structure model components from PDB IDs 7YV1, 3SOB, 7SRO, and 9NMV were docked into the map, using UCSF ChimeraX (*43*), followed by manual adjustment and rebuilding in COOT (*44*). Real-space refinement was carried out in PHENIX with secondary structure and geometry restraints applied (*45*). Model validation was performed with MolProbity, including assessment of Ramachandran statistics and overall scores (**Table S4**) (*46*). Coordinates have been deposited in the protein databank with accessions 9YTF (vAB-30/A01-EVD) and 9YTD (vAB-66/A02-PAP).

## Supporting information

designed protein sequences

## Software and Data Availability

Open source Python scripts for running the ADAPT design pipeline are freely available under an MIT license at https://github.com/phbradley/ADAPT. Fine-tuned AF2 and RFantibody parameter sets along with template and CDR3 database files can be downloaded from the Zenodo repository with DOI 10.5281/zenodo.17488258, which also contains amino acid sequences and structural models for the TCR and antibody designs.

## Acknowledgments

We acknowledge K. VanWormer, L. Goldschmidt, and Fred Hutch Scientific Computing for their outstanding lab and computational support. We benefitted from discussions with and advice from B. Vogelstein, T. Nichakawade, T. Schumacher, J. Watson, N. Bennett. We thank R. Yan, J. Jung, X. and S.Yang from the Howard Hughes Medical Institute Janelia Research Campus Cryo-EM Facility for their assistance with data collection. We thank J. Sims, M. Bauer, and M. Glögl for helping with SPR. We thank the NIH Tetramer Core Facility (NIH Contract 75N93020D00005 and RRID:SCR_026557) for providing some of the pMHC tetramers and monomers used in this manuscript.

## Author contributions

Conceptualization: PB, AM, DB; Experimental strategy: AM, DB, KCG, AMi, DF, CC, KMJ, NW; Experimental testing: AM, KMJ, NW; Software design: PB; Software testing and refinement: PB, AM, MAL; Protein design calculations: PB, AM, MAL, AE; Funding Acquisition: DB, PB, KCG, PGT; Writing – original draft: PB, AM, KMJ, NW, DB; Writing – review & editing: PB, AM, KMJ, NW, DB, AMi, KCG, PGT, MAL

