## Supplementary Data for "Targeting peptide–MHC complexes with designed T cell receptors and antibodies"

**Supplementary Table 1. Off-target peptides for flow cytometry binding experiments**

| MHC | Peptide | Off-target MHC | Off-target peptide |
|---|---|---|---|
| A*01:01 | EVDPIGHLY | A*01:01 | ESDPIVAQY |
| A*02:01 | TLMSAMTNL | A*02:01 | NLVPMVATV |
| A*02:01 | ALYDKTKRI | A*02:01 | SLLMWITQC |
| A*02:01 | GLMWLSYFV | A*02:01 | SLLMWITQC |
| A*02:01 | LLWNGPIAV | A*02:01 | SLLMWITQC |
| B*44:02 | SEITKQEKDF | A*01:01 | EVDPIGHLY |

**Supplementary Table 2. Designed sequence changes in the CDR loops**

| Loop | Number of mutations per loop mean (min - max) | Percent mutated positions per loop mean (min - max) |
|---|---|---|
| CDR1A | 5.0 (2 - 7) | 79.8% (33.3% - 100.0%) |
| CDR2A | 5.1 (3 - 8) | 76.5% (42.9% - 100.0%) |
| CDR3A[1] | 8.8 (6 - 10)[2] | 88.7% (80.0% - 100.0%) |
| CDR1B | 3.5 (2 - 5) | 70.0% (40.0% - 100.0%) |
| CDR2B | 4.4 (3 - 6) | 73.2% (50.0% - 100.0%) |
| CDR3B[1] | 7.4 (4 - 10)[2] | 82.8% (50.0% - 100.0%) |

[1]CDR3 statistics exclude the first three ('CXX') and last two (typically 'XF') positions, which are not varied in the design process.
[2]CDR3 mutation counts include indels since the length of the designed loop does not necessarily match the length of the CDR3 in the framework template.

**Supplementary Table 3. Antibody binding affinities from surface plasmon resonance.**

| Complex | Analyte | Ligand | Kd (nM) |
|---|---|---|---|
| vAB-5 : A01-EVD | monovalent pMHC | IgG | 83 |
| vAB-28 : A01-EVD | monovalent pMHC | IgG | 284 |
| vAB-30 : A01-EVD | monovalent pMHC | IgG | 656 |
| vAB-66 : A02-TLM | monovalent IgG Fab | pMHC | 151 |
| vAB-68 : A02-TLM | monovalent IgG Fab | pMHC | 157 |
| vAB-69 : A02-TLM | monovalent IgG Fab | pMHC | 409 |
| vAB-72 : A02-TLM | monovalent IgG Fab | pMHC | 520 |
| vAB-220 : A02-ALY | monovalent IgG Fab | pMHC | 747 |
| vAB-246 : A02-ALY | monovalent IgG Fab | pMHC | 393 |
| vAB-247 : A02-ALY | monovalent IgG Fab | pMHC | 5 |
| vAB-250 : A02-ALY | monovalent IgG Fab | pMHC | 23 |

**Supplementary Table 4. Summary of data collection and model statistics.**

| | vAB-30/A01-EVD | vAB-66/A02-TLM |
|---|---|---|
| EM equipment | FEI Titan Krios | FEI Titan Krios |
| Voltage (kV) | 300 | 300 |
| Detector | K3 | Falcon 4i |
| Pixel size (Å) | 0.827 | 0.94 |
| Electron dose ($e^-$/Å$^2$) | 60 | 50 |
| Defocus range (µm) | (-1.0, -2.0) | (-0.8, -2.0) |
| **Reconstruction** | | |
| Software | CryoSPARC | CryoSPARC |
| Number of used Particles | 1,317,109 | 880,830 |
| Map sharpening Method | deepEMhancer (visualization) | Uniform |
| Final Resolution (Å) | 2.6 | 2.5 |
| **Model building and refinement** | | |
| Software | PHENIX & COOT | PHENIX & COOT |
| Initial models used (PDB codes) | Predicted structure | 3SOB, 7YV1, 7SR0, 9NMV |
| Model composition | | |
| Non-hydrogen atoms | 7270 | 7368 |
| Protein residues | 932 | 933 |
| Water molecules | 0 | 64 |
| B factors (Å$^2$) | | |
| Protein | 47.37 | 39.28 |
| Water | —— | 35.36 |
| R.m.s deviations | | |
| Bonds length (Å) | 0.004 | 0.003 |
| Bonds Angle (˚) | 0.503 | 0.548 |
| Ramachandran plot statistics (%) | | |
| Preferred | 98.04 | 98.70 |
| Allowed | 1.96 | 1.30 |
| Outlier | 0.00 | 0.00 |
| Validation | | |
| Molprobity score | 1.24 | 1.17 |
| Clash score | 4.62 | 2.51 |
| Rotamer outliers (%) | 0.00 | 1.49 |

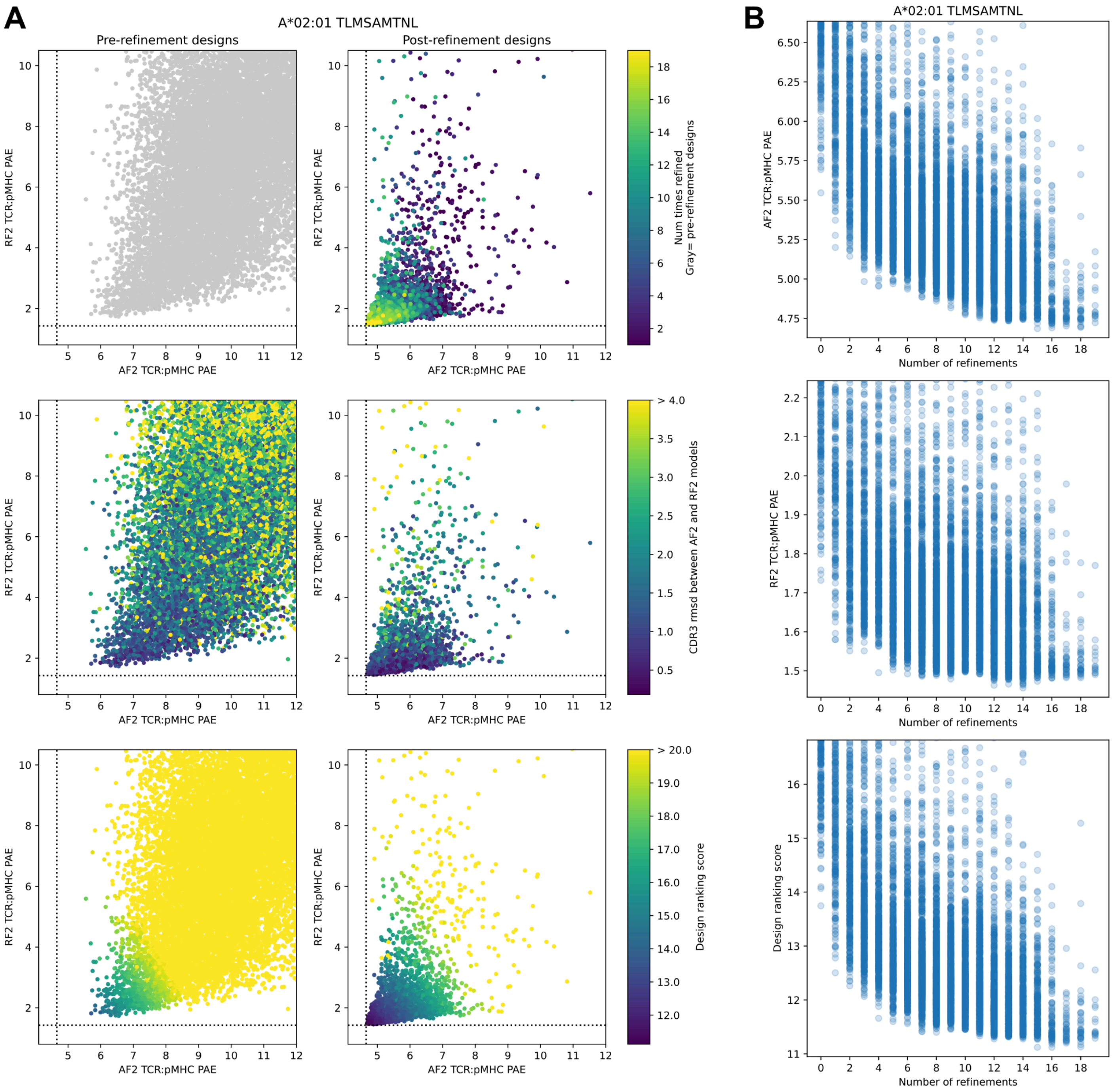


**Supplementary Figure 1. Design quality metrics before and after refinement. (A)** Scatter plots of AF2 predicted aligned error (PAE) between TCR and pMHC (x-axis) versus RF2 PAE between TCR and pMHC (y-axis) for pre- and post-refinement designs (left and right columns, respectively). Each dot represents an individual design. Scatter plots are colored by the number of refinement cycles (top row), the CDR3 RMSD between the AF2 and RF2 models of the design (middle row, calculated after superimposing on the MHC), and the design ranking score (bottom row), which is a weighted combination of the two inter-PAE values and the CDR3 RMSD (see Methods). **(B)** Scatter plots of design metrics (y-axis) as a function of the number of refinement cycles (x-axis); pre-refinement designs are plotted at x=0.

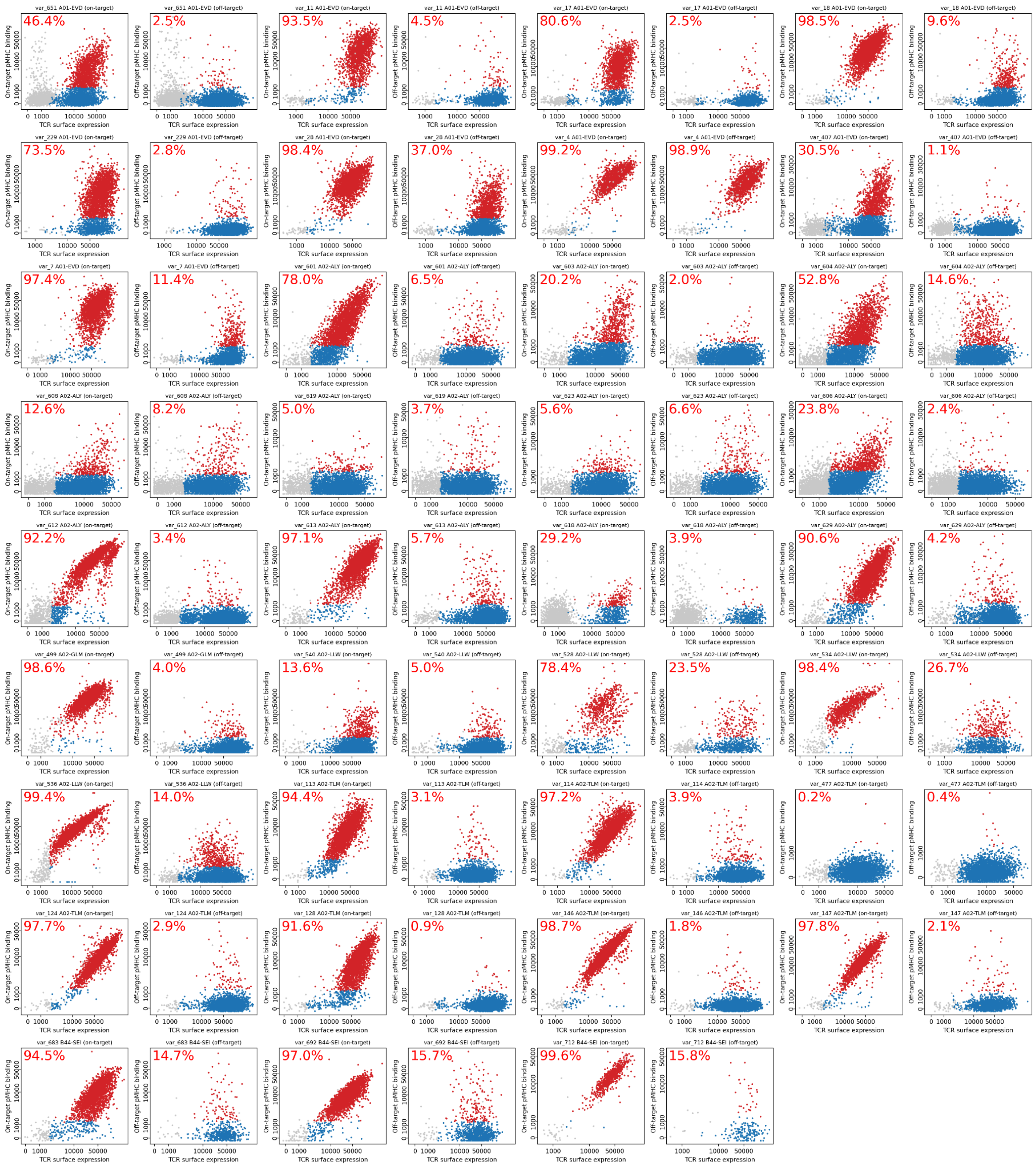


**Supplementary Figure 2. Representative pMHC binding data for 35 designed TCRs analyzed by flow cytometry.** TCR surface expression as measured by CD3 staining is plotted on the x-axis; on- or off-target binding to multimerized pMHC is plotted on the y-axis. Cells gated as TCR-positive are shown in blue (if gated as pMHC-negative) or red (if gated as pMHC-positive). Labels in the upper left corner represent the percentage of TCR-positive cells that are pMHC positive.

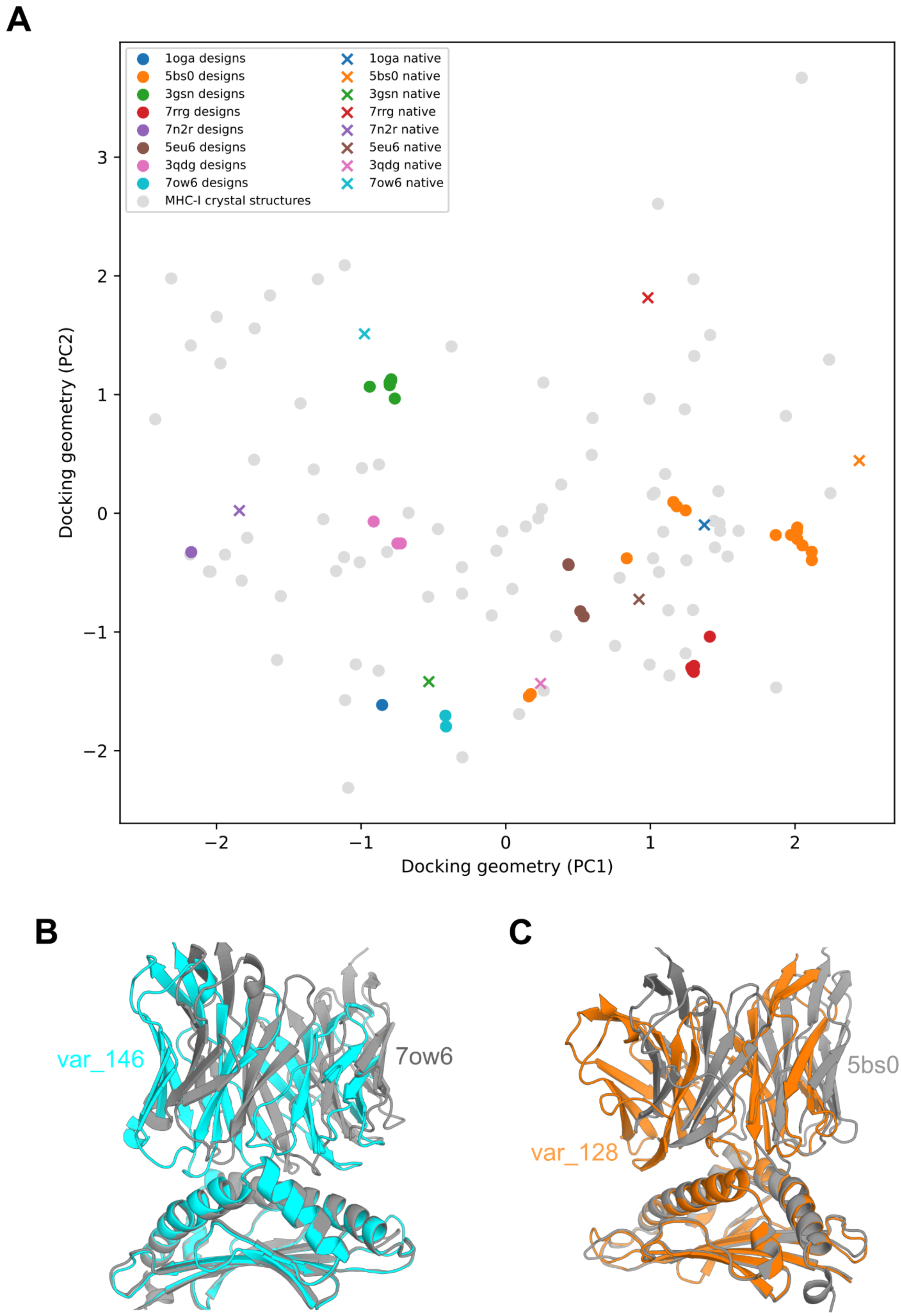


**Supplementary Figure 3. Comparison of docking geometries in design models and their framework-region template structures.** (**A**) Scatter plot of the top two of 6 principal components of the TCR:pMHC rigid-body docking space for binding modes from a set of successful design models (disks colored by the native structural template for their framework region) and a set of native class I TCR:pMHC structures (gray disks and colored Xs for those template TCRs). Designed binding modes are not simply borrowed from the framework template, as expected since only the internal coordinates of the template TCR, and not the docking geometry, are supplied to AF2 during ADAPT design. **(B-C)** Example superpositions of design models (cyan and orange) and framework template structures (gray), aligned on the MHC chain.

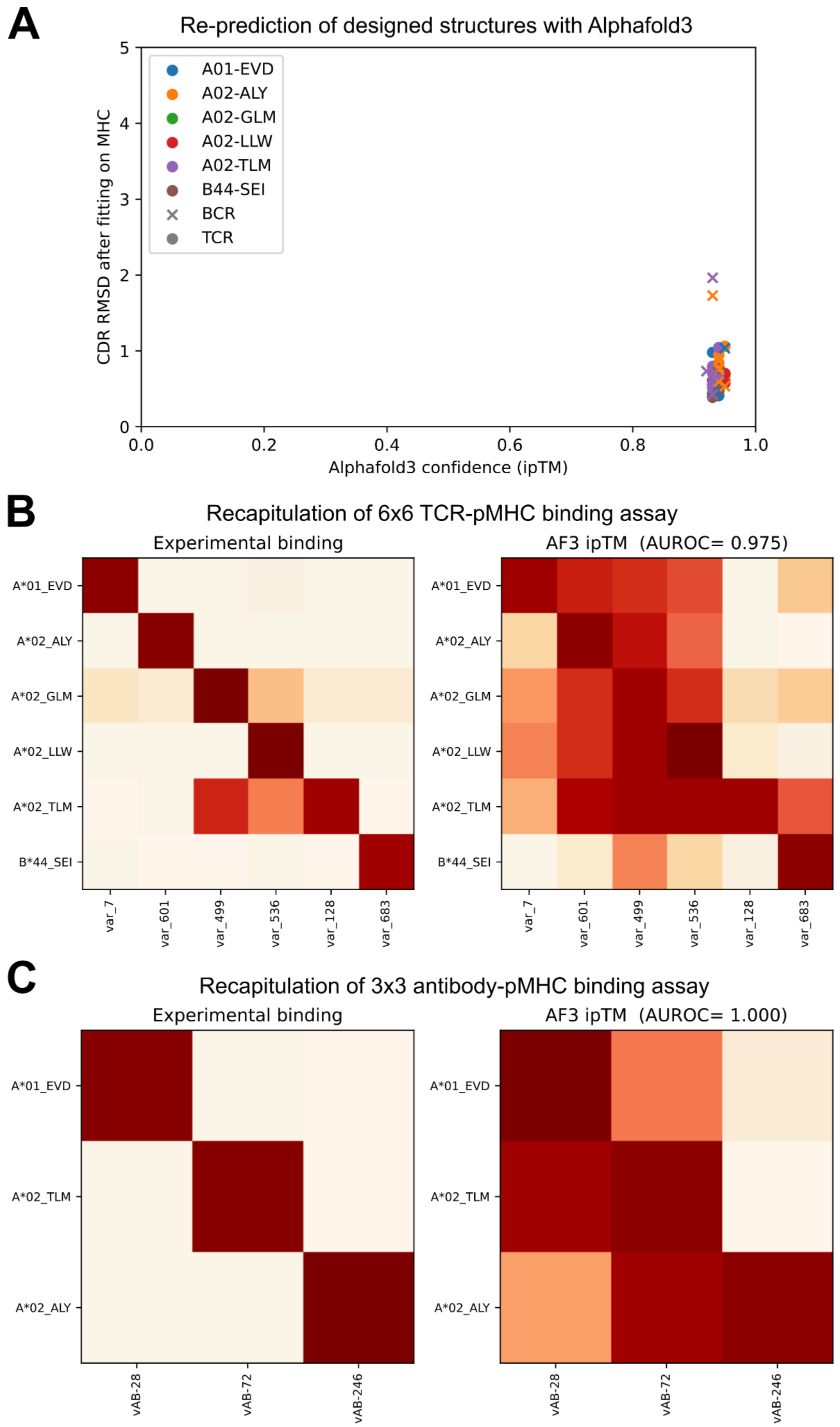


**Supplementary Figure 4. Retrospective analysis of TCR and antibody designs with Alphafold3. (A)** The default Alphafold3 pipeline was used to predict the designed interface structures using as input only the designed sequences (without interface templates or fine-tuning). Ca-RMSD values calculated over the CDR loops after aligning on MHC (y-axis) were below 1Å for nearly all the designs, and interface confidence was high as indicated by ipTM scores (x-axis) consistently above 0.9. **(B)** Heatmaps of experimental binding (left) and AF3 ipTM scores (right) for all 6x6=36 pairings of the 6 pMHC targets and a single representative TCR design for each. Ranking the possible pairings by ipTM and considering the designed interactions (diagonal entries) as true interactions gives an area under the receiver operating characteristic curve (AUROC) value of 0.975. **(C)** Experimental (left) and AF3-simulated (right) cross-binding heatmaps for three pMHC targets and a single representative antibody design for each. Ranking the possible pairings by AF3 ipTM score gives an AUROC of 1.0 for discriminating the designed pairings (i.e., the diagonal of the heatmap) from off-target interactions (off-diagonal entries in the heatmap).

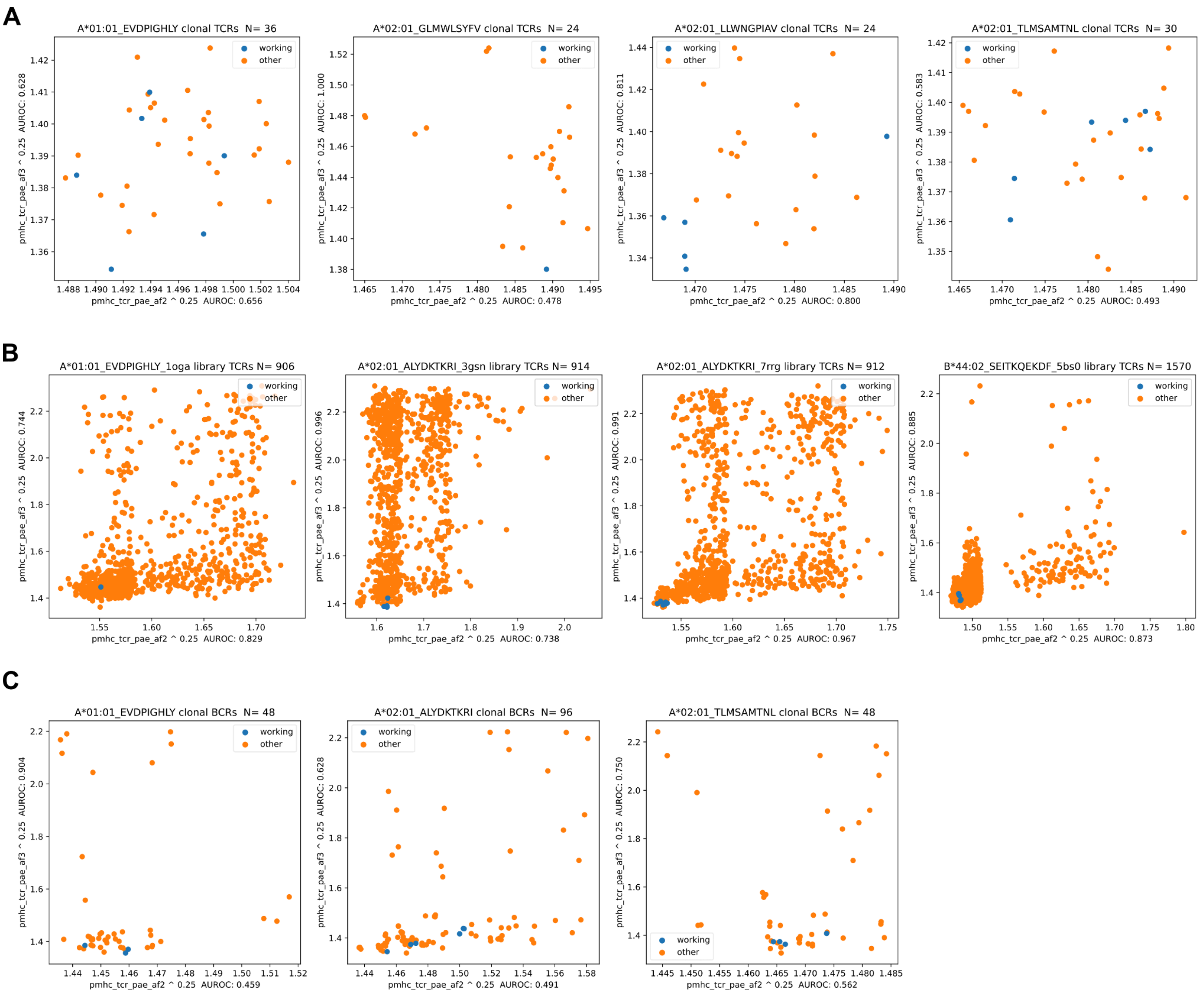


**Supplementary Figure 5. Comparison of Alphafold interface metrics for working and non-working designs.** Scatter plots of Alphafold2 (x-axis) versus Alphafold3 (y-axis) TCR-pMHC interface PAE scores for working and non-working designs; lower values indicate lower predicted error and hence higher confidence in the designed interface. Raw PAE values are raised to the 1/4th power to focus on the lowest scores and compress the higher scores corresponding to low-confidence designs. Area under the receiver operating characteristic curve (AUROC) values measuring discrimination of working from non-working designs are provided in the corresponding axis labels. Discrimination of working designs is modest for clonal TCR designs **(A)** but stronger for TCR libraries **(B)** and antibodies **(C)**.

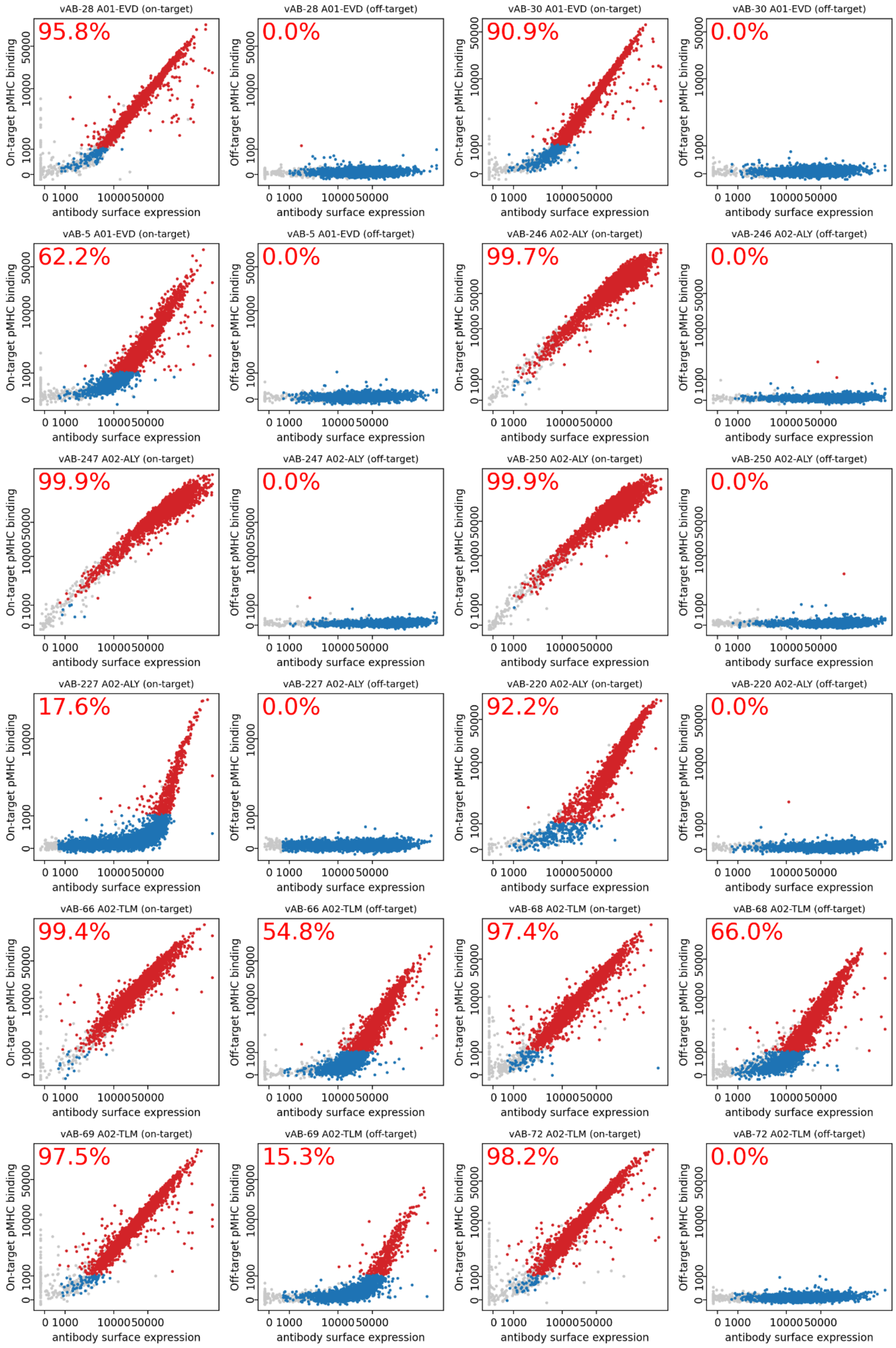


**Supplementary Figure 6. Representative pMHC binding data for 12 designed antibodies analyzed by flow cytometry.** IgG surface expression is plotted on the x-axis; on- or off-target binding to multimerized pMHC is plotted on the y-axis. Cells gated as IgG-positive are shown in blue (if gated as pMHC-negative) or red (if gated as pMHC-positive). Labels in the upper left corner represent the percentage of IgG-positive cells that are pMHC positive.

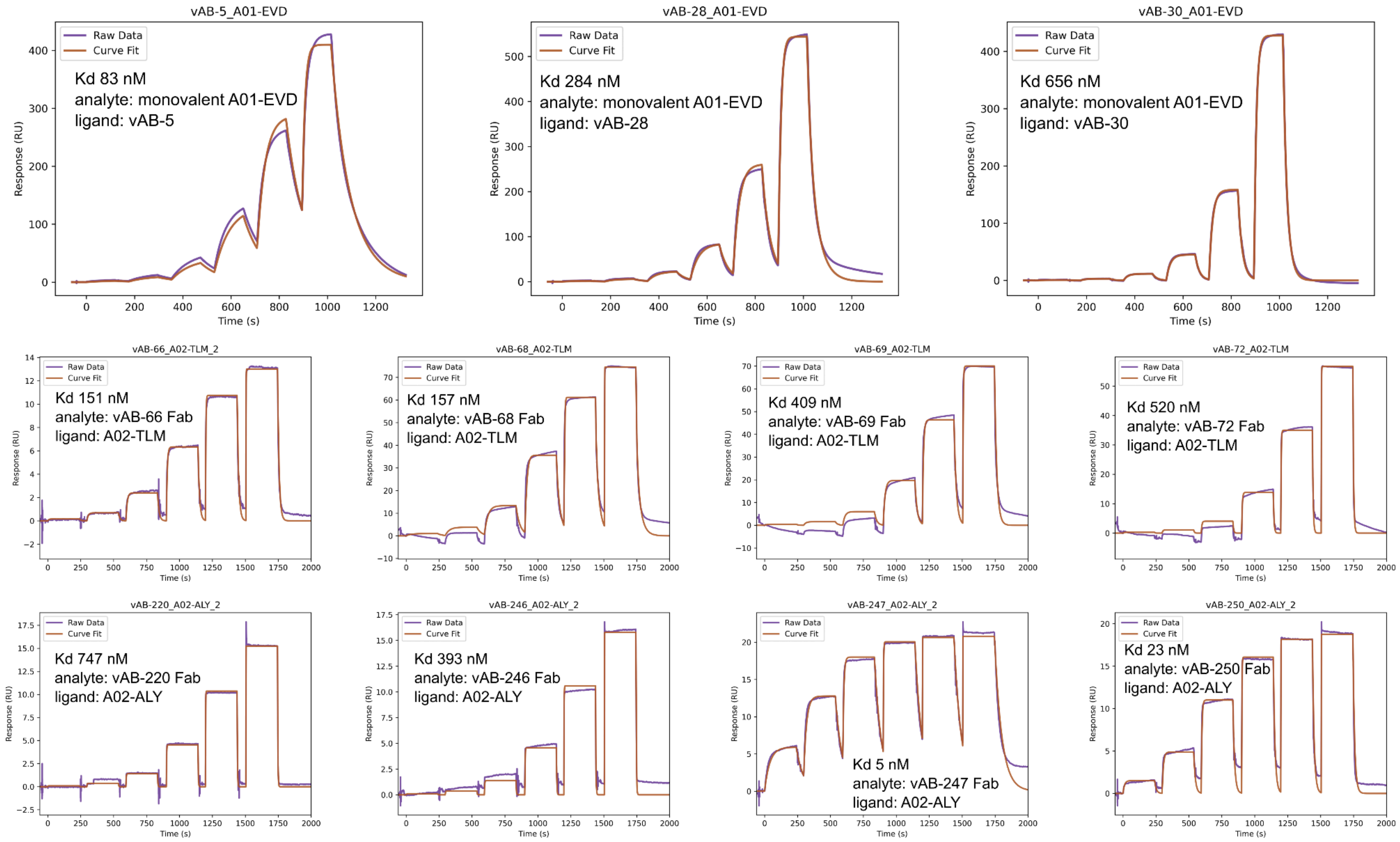


**Supplementary Figure 7. Surface plasmon resonance (SPR) analysis of designed antibody–pMHC interactions.** Binding responses were measured on a Biacore 8K at 25 °C in HBS-EP+ buffer. For the A01-EVD system (top row), IgG1s were captured on a Protein A chip and monovalent biotinylated A01-EVD analytes were injected. For all other targets (A02-TLM, A02-ALY), biotinylated pMHCs were immobilized on a streptavidin chip and IgG1-Fab analytes were tested using single-cycle kinetics. Raw sensorgrams (purple) and global 1:1 Langmuir fits (maroon) are shown with dissociation constants ($K_d$) indicated.

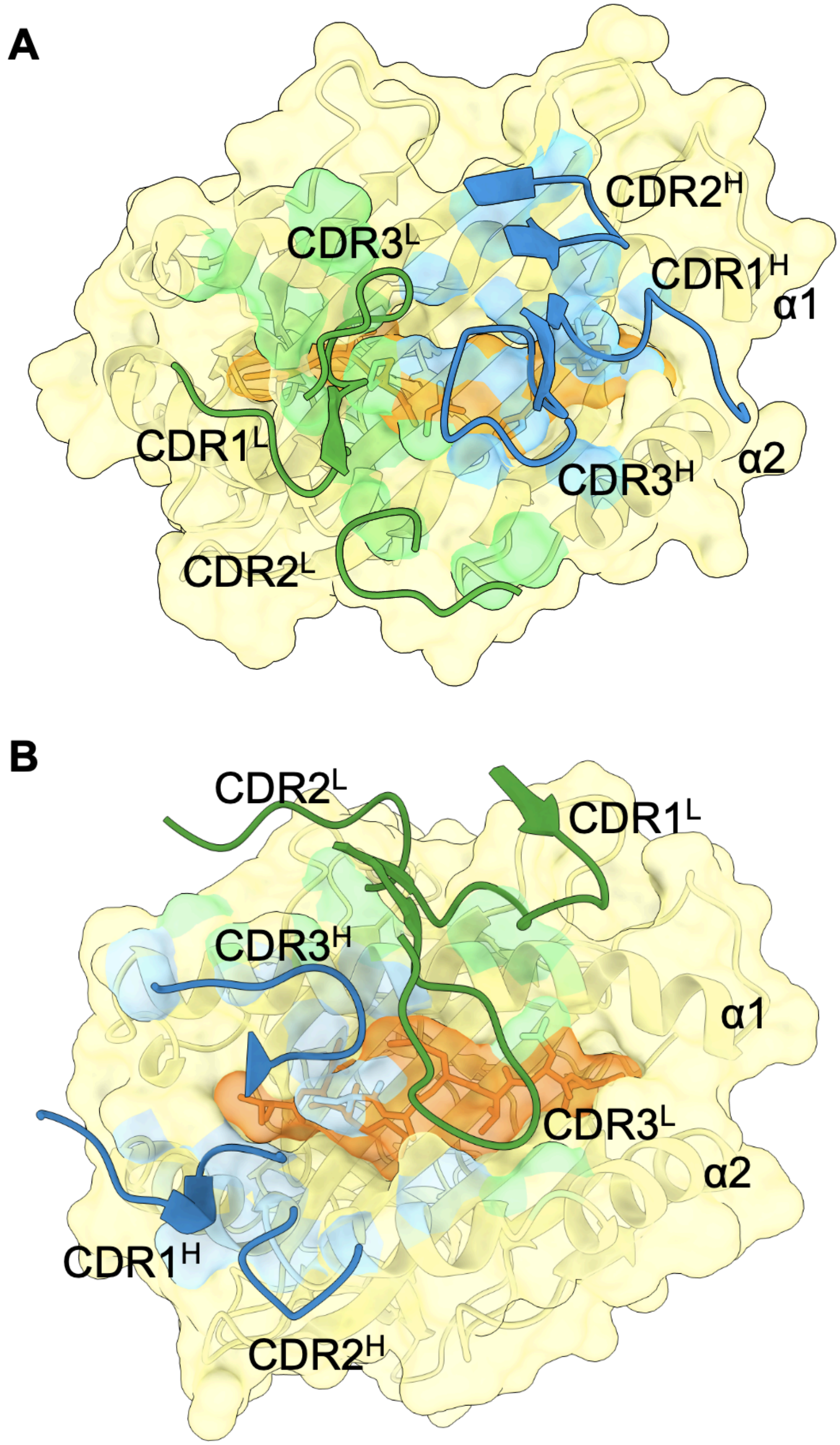


**Supplementary Figure 8. CDR loop interactions with pMHC in cryoEM structures.** MHC is shown in pale yellow, peptide in orange, Fab heavy chains in blue, and Fab light chains in green. Contacts to pMHC from the heavy chain are shown as light blue and from the light chain are shown as light green. (A) vAB-30/A01-EVD (B) vAB-66/A02-TLM.

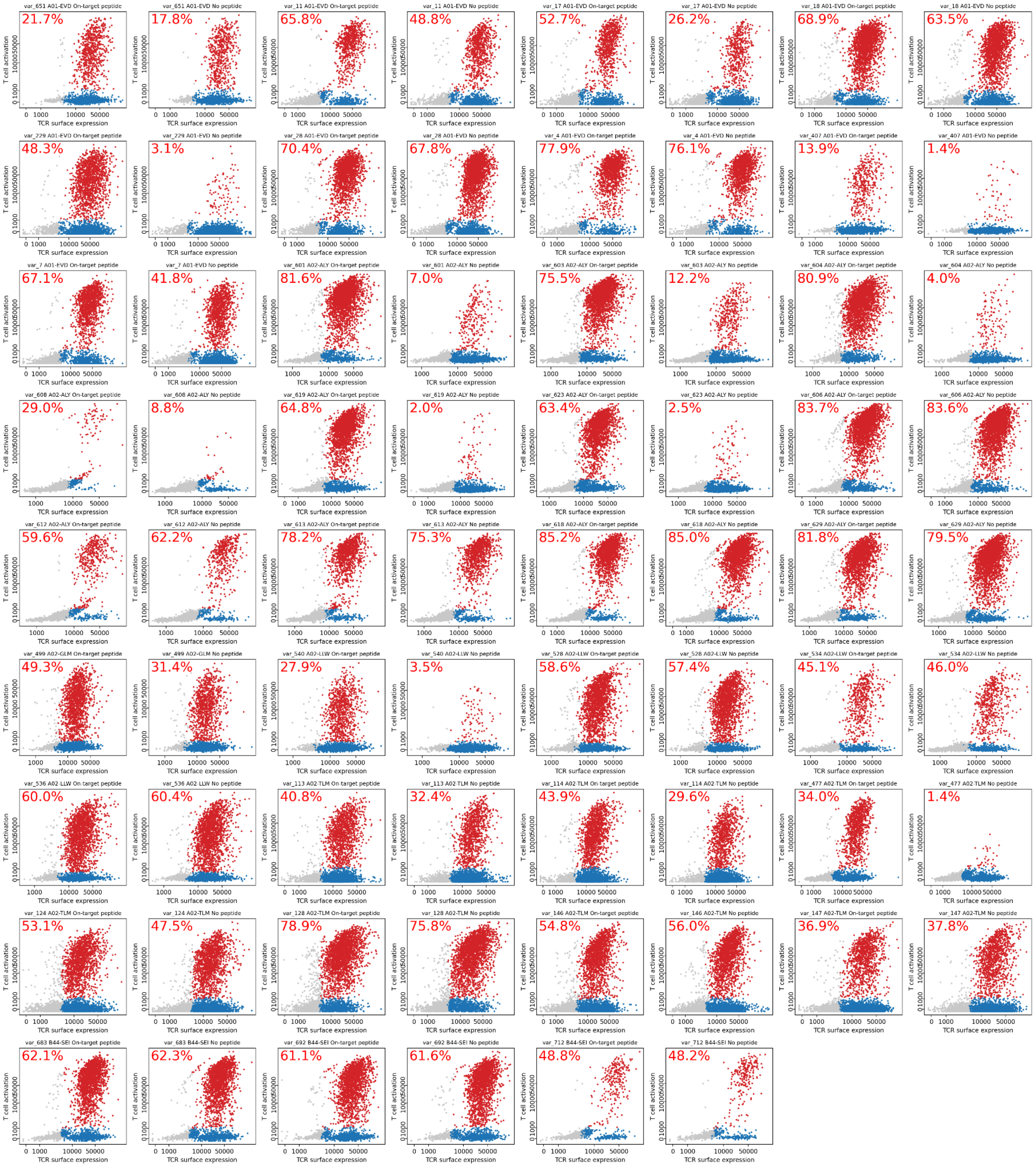


**Supplementary Figure 9. Representative T cell activation data for 35 designed TCRs analyzed by flow cytometry.** TCR surface expression as measured by CD3 staining is plotted on the x-axis; T cell activation (NFAT RE eGFP expression) is plotted on the y-axis. Cells gated as TCR-positive are shown in red if gated as active or blue otherwise. Labels in the upper left corner represent the percentage of TCR-positive cells that are activated. The two panels for each design represent activation in the presence or absence of pulsed target peptide.

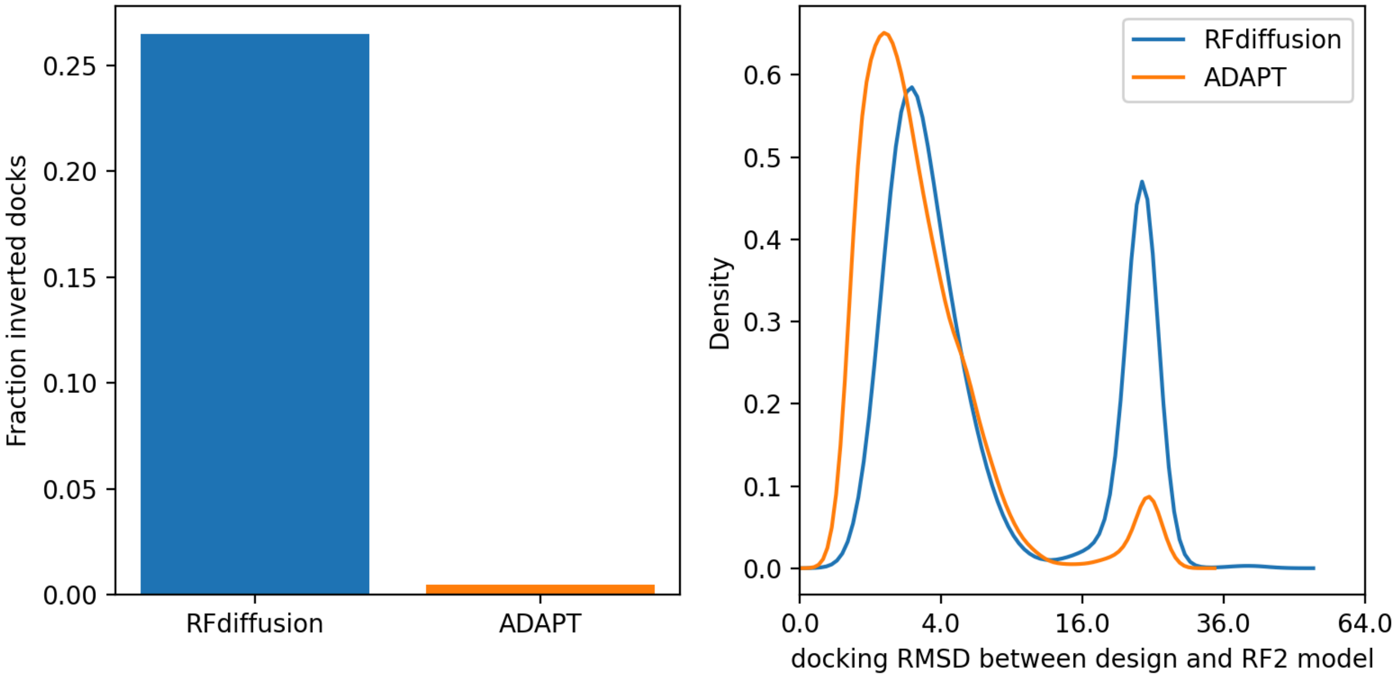


**Supplementary Figure 10. Comparison between RFdiffusion- and ADAPT-generated TCR:pMHC binding modes.** A version of RFdiffusion that was fine-tuned on native TCR:pMHC complexes generated reversed binding modes at higher frequencies than the ADAPT pipeline (left panel). Reprediction of designed binding modes by RosettaFold2 was less successful for RFdiffusion designs than for ADAPT-generated designs as measured by higher docking RMSD values between the design model and the RF2 prediction (right panel).

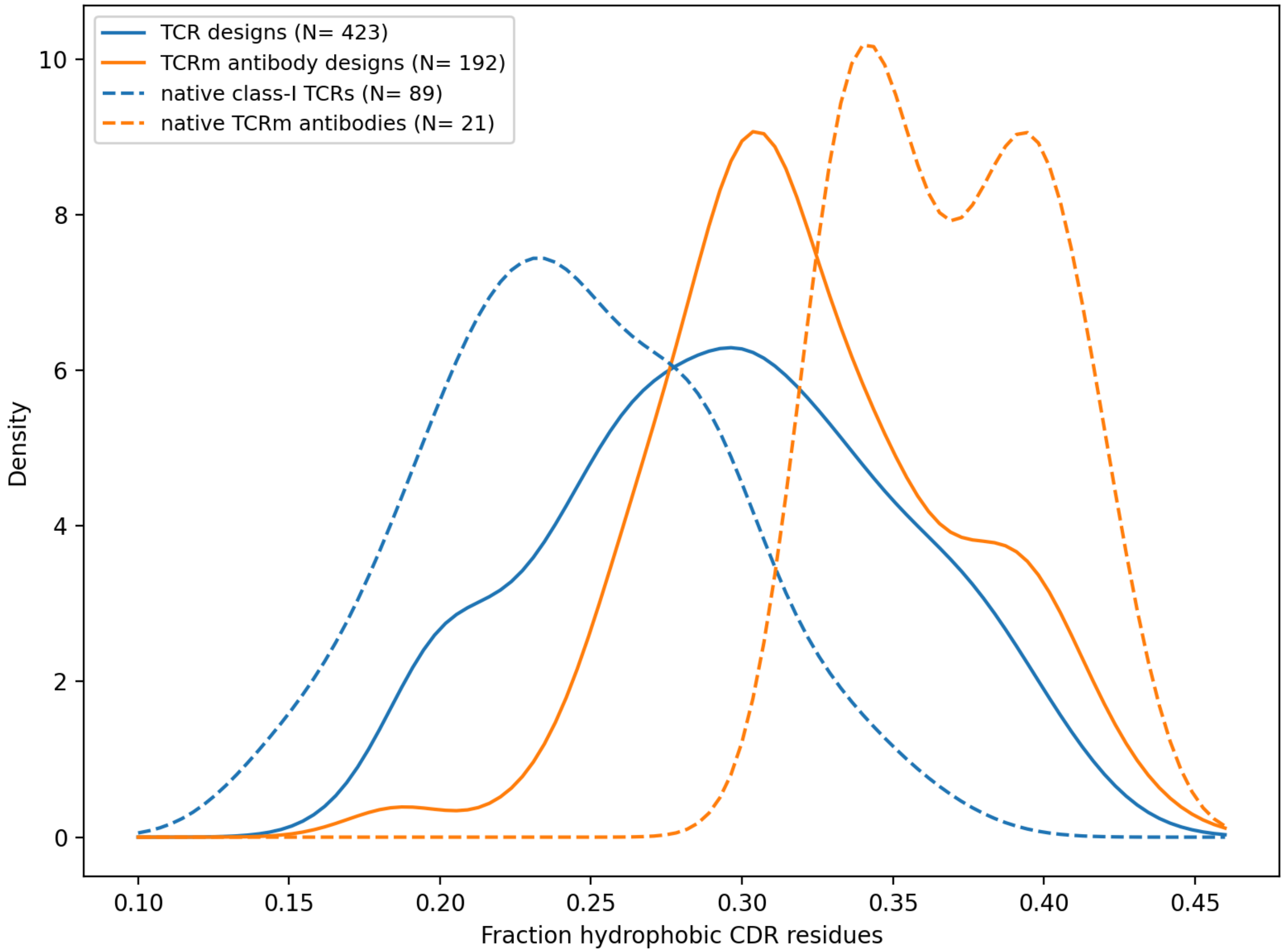


**Supplementary Figure 11. The CDR loops of designed TCRs and TCR-mimic antibodies are more hydrophobic than those of native TCRs.** Smoothed kernel density estimates of CDR loop hydrophobicity as measured by the fraction of CDR loop positions occupied by the amino acids {V,I,L,M,F,W,Y,C}. Interestingly, native TCR-mimic antibodies show even greater CDR loop hydrophobicity (orange dashed line).

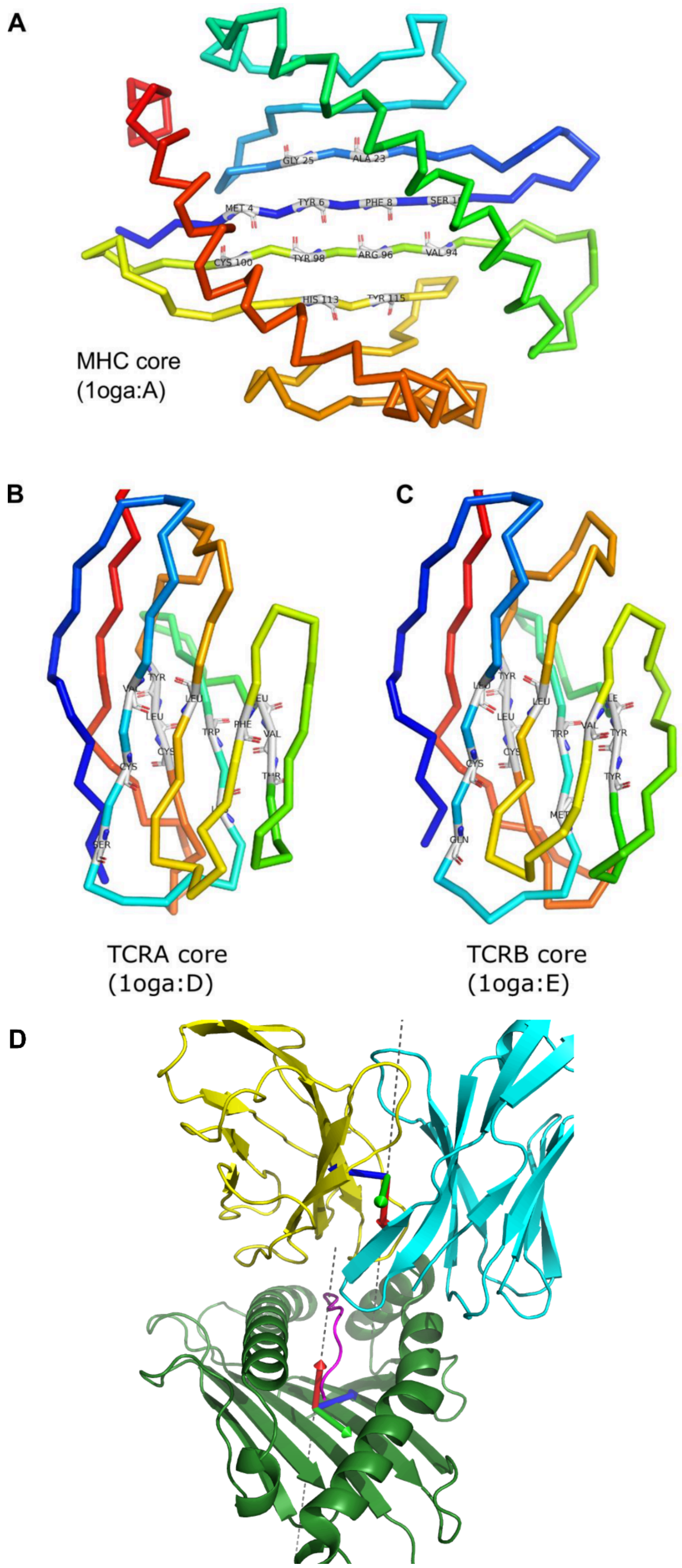


**Supplementary Figure 12. Construction of reference coordinate frames for MHC and TCR structures.** As described in the Methods, the reference frames are defined by the approximate 2-fold symmetry axis relating a set of corresponding residue pairs in the MHC beta sheet **(A)** and in the TCR alpha and beta chain framework regions **(B-C)**. **(D)** The frame origin is located at the center of mass of the aligned residues. The x-axis (red arrow) is parallel to the symmetry axis, and the z-axis (blue arrow) is directed between the centers of mass of the individual residue subsets (ie, from the N-terminal to C-terminal MHC beta-sheet halves and from the TCR alpha to beta chains).

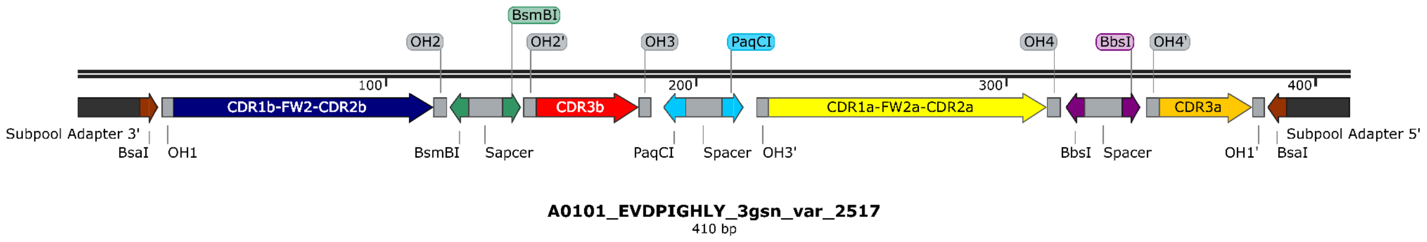


**Supplementary Figure 13. Schematic layout of pooled fragment design.** CDR1b–FW2–CDR2b, CDR3b, CDR1a–FW2a–CDR2a, and CDR3a sequences corresponding to the designed TCRs with a specified framework were interspersed with golden gate assembly sites to enable sequential cloning for integrating the invariant intermediating DNA fragments as described in Methods.

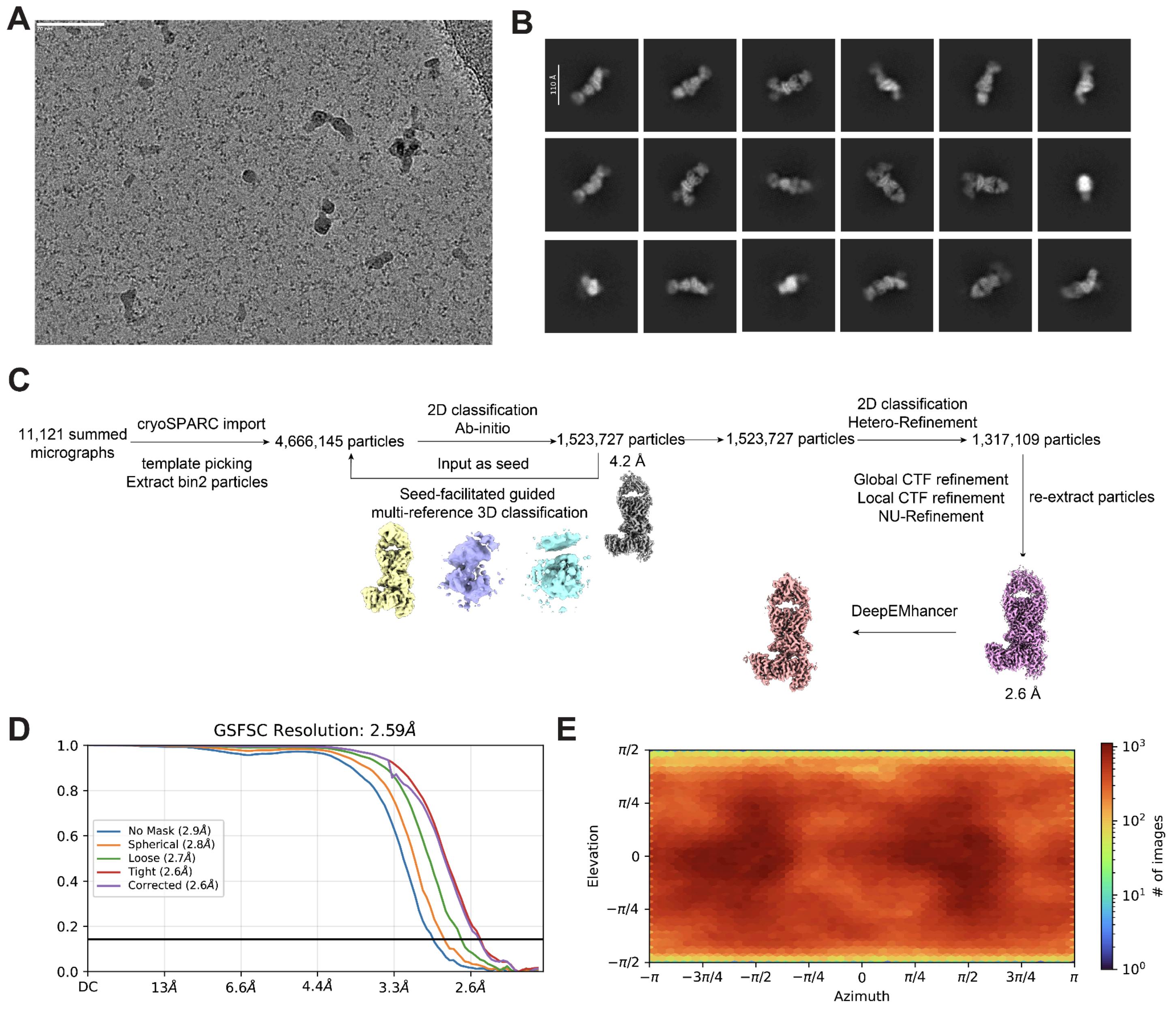


**Supplementary Figure 14. Workflow for cryo-EM data processing for vAB-30/A01-EVD complex.** (**A**) A representative micrograph of the vAB-30/A01-EVD complex. Scale bar = 70 nm. (**B**) Representative 2D class averages illustrating the varied orientations of the vAB-30/A01-EVD. (**C**) Flow chart for data processing of the vAB-30/A01-EVD complex. (**D**) Gold standard FSC curve illustrating the resolution determination for the vAB-30/A01-EVD complex, achieved through iterative 3D refinement. The resolution at the FSC 0.143 criterion is 2.6 Å. (**E**) Angular distribution of the particles used for the final reconstructions.

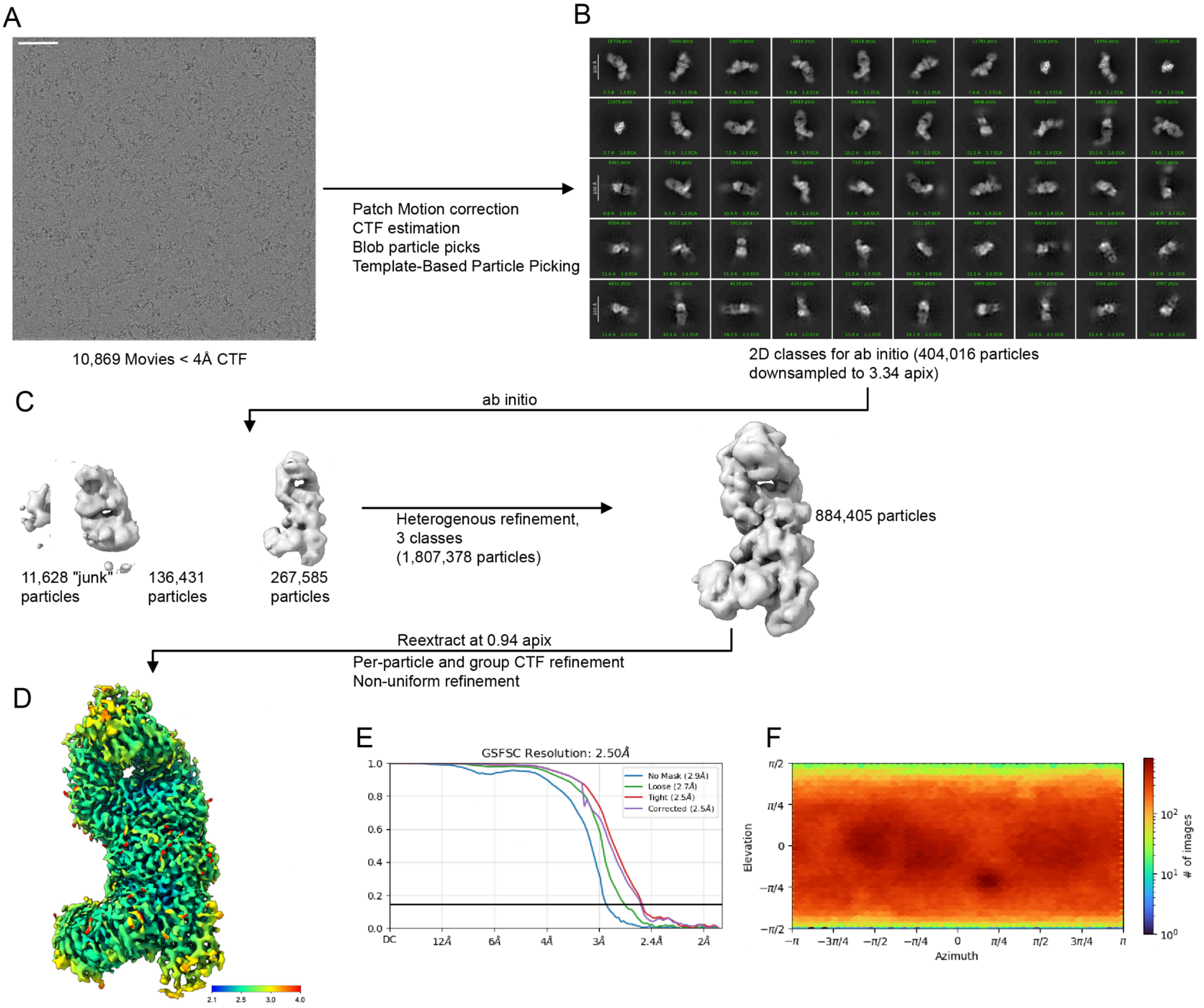


**Supplementary Figure 15. Workflow for cryo-EM data processing for vAB-66/A02-TLM complex.** (**A**) A representative micrograph of the vAB-66/A02-TLM complex. Scale bar = 50 nm. (**B**) Representative 2D class averages illustrating the varied orientations of the vAB-66/A02-TLM. (**C**) Flow chart for data processing of the vAB-66/A02-TLM complex. (**D)** Final map colored by local resolution. (**E**) Gold standard FSC curve illustrating the resolution determination for the vAB-66/A02-TLM complex, achieved through iterative 3D refinement. The resolution at the FSC 0.143 criterion is 2.5 Å. (**F**) Angular distribution of the particles used for the final reconstructions.
